# Repeatable, low-drift recordings in behaving non-human primates using flexible microelectrodes

**DOI:** 10.64898/2026.01.09.698500

**Authors:** Daniel P. Woods, Grace M. Adams, Rana Mozumder, Wenhao Dang, Andrew Y. Chen, Christos Constantinidis, Daniel L. Gonzales

## Abstract

Neurophysiological recordings from non-human primates (NHPs) have traditionally relied on rigid microelectrode arrays made from stainless steel or silicon. While these devices enable high-quality recordings, a fundamental mechanical mismatch between rigid materials and soft brain tissue leads to inflammation, gliosis, and signal instability. In particular, brain micromotion causes continuous drifting of neurons relative to fixed electrodes, compromising single-unit tracking during both chronic and acute recordings. Flexible, penetrating electrodes offer a promising solution, but their adoption in NHPs has been hindered by the technical challenges of delivering ultra-thin polymers through thick dura mater. Here, we demonstrate a comprehensive approach for acute, repeated recordings in awake, behaving NHPs using flexible arrays. We fabricated a microelectrode array that spans cortical layers with 32 cellular-scale recording sites embedded in 7 μm-thick Parylene-C. We developed a novel “telescopic” insertion method that combines concentric guide tubes with a retractable microwire shuttle. Our technique is compatible with standard chronic recording chambers and allowed for repeated penetration of free-floating arrays through intact dura over weeks without the need for a new craniotomy. Across two awake rhesus macaques, we optimized the electrode geometry and insertion procedure to achieve an 80% single-unit recording success rate. As animals performed an oculomotor delayed response task, we recorded task-responsive neurons from prefrontal and posterior parietal cortex with stable single-unit activity throughout 1–2-hour behavioral sessions. Critically, by comparing our flexible arrays to rigid probes in the same animals and recording chambers, we provide quantitative evidence that flexible electrodes reduce total single-unit drift from hundreds to tens of microns. Our work establishes flexible microelectrode arrays as a practical, dependable technology for NHP neuroscience and paves the way toward long-term, ultra-stable neurophysiology in large animal models.

## INTRODUCTION

High-fidelity neurophysiological recordings obtained from non-human primates (NHPs) are essential for mapping the circuit dynamics of cognition, yet current technologies face fundamental limitations that compromise recording quality and long-term stability. Multi-site recordings in these large-brained animals have traditionally relied on devices constructed from rigid materials. For example, steel microelectrode arrays such as Plexon S/U/V-probes have been a staple of NHP neurophysiology for decades (Bastos et al., 2018; Pagan et al., 2013) and silicon Utah Arrays the technological basis for high performing brain-computer interfaces (Carmena et al., 2003; Oby et al., 2019; Sadtler et al., 2014; Santhanam et al., 2006). More recently, high-density silicon probes such as Neuropixels have revolutionized the field by recording from hundreds of neurons simultaneously with unprecedented spatial resolution (Mozumder et al., 2026; Trautmann et al., 2025). Despite these technological advances, all rigid electrode technologies share a fundamental limitation: a mechanical mismatch between stiff materials (Young’s moduli ∼200 GPa for steel and silicon) and soft brain tissue (∼1-10 kPa) (He et al., 2020; Thielen & Meng, 2021). This orders-of-magnitude difference in stiffness leads to significant complications at the tissue-electrode interface, including chronic inflammation, glial scarring, and progressive signal instability over time (Kozai et al., 2015; Salatino et al., 2017). Brain micromotion due to respiration, cardiac rhythm, and cerebrospinal fluid (CSF) flow exacerbates the challenges faced by rigid probes (Forrest et al., 2025; Gilletti & Muthuswamy, 2006). This motion causes shearing against the inserted neural interface, resulting in the physiological drift of neurons relative to the fixed position of microelectrodes. In large animals and humans, this motion can be on the order of hundreds of microns (Chung et al., 2022; Paulk et al., 2022). Even during acute recordings lasting only a few hours, drift compromises the ability to track individual neural units and complicates data analysis (Boussard et al., 2023; Pachitariu et al., 2021). For chronic implants, drift leads to large fluctuations in the day-to-day yield of unit activity (Hahn et al., 2025).

To mitigate these challenges, flexible penetrating electrodes have emerged as a promising alternative to rigid probes. Materials such as polyimide, Parylene-C, and SU-8 provide enhanced mechanical compliance with brain tissue (Song et al., 2020). In rodents, there is robust evidence that flexible thin-film electrodes integrate with neural tissue, reduce gliosis and scarring, and provide stable recordings during long-term implantation (Chung, Joo, Fan, et al., 2019; Kim et al., 2013; Luan et al., 2017; Musk & Neuralink, 2019; Yasar et al., 2024; S. Zhao et al., 2023; Z. Zhao et al., 2022; Zhou et al., 2017). However, these technologies have been slow to be adopted for NHPs. The challenge of neurophysiology is magnified in large animals by intensive, high-risk surgeries and thick dura matter that must be removed or penetrated for intracranial recordings. Surgical removal of the dura carries risks such as potential CSF leakage, inflammation, and infection (Overton et al., 2017). Dura penetration is not without risk as well. Brittle probes can break or become damaged during insertion. Even rigid electrodes such as silicon Neuropixels and steel Plexon probes often require large “guide tubes” that penetrate through dura and allow for microelectrode array insertion (Namima et al., 2024; Trautmann et al., 2025). Flexible electrodes face an even greater challenge. Their mechanical compliance—the very property that makes them appealing for tissue integration—renders them difficult to deploy as free-floating arrays in brain tissue. Current insertion methods for flexible electrodes require complex, customized combinations of temporary stiffening shuttles and biodegradable coatings, often in conjunction with complete dura removal during open skull surgical procedures (Chung, Joo, Smyth, et al., 2019; Thielen & Meng, 2021; Z. Zhao et al., 2019). Compared to well-established techniques in NHPs for the repeated insertion of rigid arrays through chronic recording chambers, this additional complexity has made the practical advantages of flexible electrodes unclear. Therefore, while flexible electrodes theoretically offer a new frontier of low-footprint, highly stable recordings in NHPs, the field lacks clear demonstrations that the benefits outweigh the substantial challenges.

Existing studies using flexible neural interfaces in NHPs indeed suggest that this approach offers a stable interface to the brain (Gerbella et al., 2021; Jeanpierre et al., 2025; K. Lee et al., 2024; Y. Liu et al., 2024; Merken et al., 2022; Oh et al., 2025; Pothof et al., 2016; Qiang et al., 2025; Tian et al., 2023; Y. Wang et al., 2023). Chronic polyimide implants have been able to record local field potentials (Oh et al., 2025) and spiking activity (Gerbella et al., 2021; Merken et al., 2022; Tian et al., 2023), with some instances of reliable single-unit tracking across recording sessions (Tian et al., 2023). Proof-of-concept recordings with novel flexible geometries and materials have also been performed in NHPs in acute settings, typically in anesthetized animals (Jeanpierre et al., 2025; K. Lee et al., 2024; Y. Liu et al., 2024; Qiang et al., 2025). In the vast majority of both chronic and acute cases, these implantations were conducted following a new craniotomy and dura removal. In only two studies, flexible microelectrode arrays have been inserted through a traditional chronic recording chamber without the need for a new craniotomy (Y. Liu et al., 2024; Y. Wang et al., 2023). In both instances, the stiffening shuttled used for insertion was either not removed or unable to be reliably removed, therefore impeding the full potential of the flexible array. Together, these studies highlight the promise of emerging capabilities—including wireless recording (Oh et al., 2025) and increasingly large-scale, high-density flexible arrays (Y. Liu et al., 2024; Tian et al., 2023)—but also underscore that flexible neural interfaces in NHPs remain a nascent and technically constrained field.

We identify two key areas that need further development to usher in an era of ultra-stable, single-neuron recordings in NHPs with flexible arrays. First, we must establish robust, low-risk implantation protocols that can be easily adopted by NHP research groups. This requires approaches that circumvent the need for a new craniotomy prior to every session and have a high likelihood of successful recording. While several insertion techniques have been demonstrated for NHPs (Y. Liu et al., 2024; Merken et al., 2022; Oh et al., 2025; Tian et al., 2023; Y. Wang et al., 2023), studies typically report only a few recordings and rarely provide statistics regarding the number of recordings attempts and successes. Without quantitative data on these surgical workflows, the barrier to entry for NHP neurophysiology groups remains high. Second, we need direct, quantitative demonstrations comparing the recording stability of rigid and flexible microelectrodes. While flexible probes are hypothesized to reduce drift by complying with brain micromotion, there are no side-by-side comparisons between flexible and rigid arrays to validate this. The current lack of data in these two areas—reliable insertion and validated stability—obscures the advantages of flexible electrodes compared to established technology and hinders their adoption by the broader NHP community.

Here, we present a comprehensive approach to these challenges by detailing a system for non-surgical, repeatable, and highly stable single-unit recordings in behaving NHPs using flexible microelectrodes. We introduce a “telescopic insertion” method that combines a conventional guide tube with a retractable shuttle mechanism for deploying free-floating arrays through intact dura. Our non-surgical insertion occurs through standard chronic recording chambers in awake animals without the need for a new craniotomy and dura resection, allowing for daily recording sessions for multiple weeks. This approach also mirrors conventional methods to NHP recordings, therefore lowering the barrier for adoption. We performed acute recordings in prefrontal cortex (PFC) and posterior parietal cortex (PPC) in actively behaving NHPs performing a working memory task. We report 21 total recordings, far surpassing similar studies with flexible arrays. After refining our electrodes and methodology—we achieved a recording success rate in 80% of sessions. Furthermore, in 18 recordings we also tracked animals behavior and observed high-yield, high signal-to-noise single-units that were associated with task-relevant variables. Importantly, our implantation approach allowed us to perform the first systematic quantification of the stability and drift associated with free-floating flexible probes in the NHP cortex. When compared to rigid microelectrode arrays, we found that flexible electrodes dramatically reduced the drift of single-unit recordings. By answering the critical insertion and stability questions that have—until now—hindered their translation to large animal models, our research paves the way for flexible neural interfaces in NHPs that are reliable, chronic, and longitudinally track individual neurons.

## RESULTS

### Fabrication of mechanically stable flexible arrays

To enable both the flexibility needed for tissue integration and the mechanical robustness required for insertion through intact dura, we designed custom Parylene-C arrays that strategically balance these competing demands (Fig. 1). We fabricated flexible arrays using a traditional combination of chemical vapor deposition of Parylene-C, photolithography, and thin-film sputter deposition (Fig. S1, Methods) (Ortigoza-Diaz et al., 2018). Probes consisted of a 32-channel linear array spanning approximately 1 mm (Fig. 1A). We specifically designed and evaluated two sets of Pt microelectrode arrays, one with a checkerboard distribution of electrodes, and the other with a linear array at the probe edge. Each recording site had a diameter of 30 µm, yielding a ∼706 µm² contact area and mean electrochemical impedance at 1 KHz of approximately 940 kΩ (Fig 1B).

**Figure 1.**
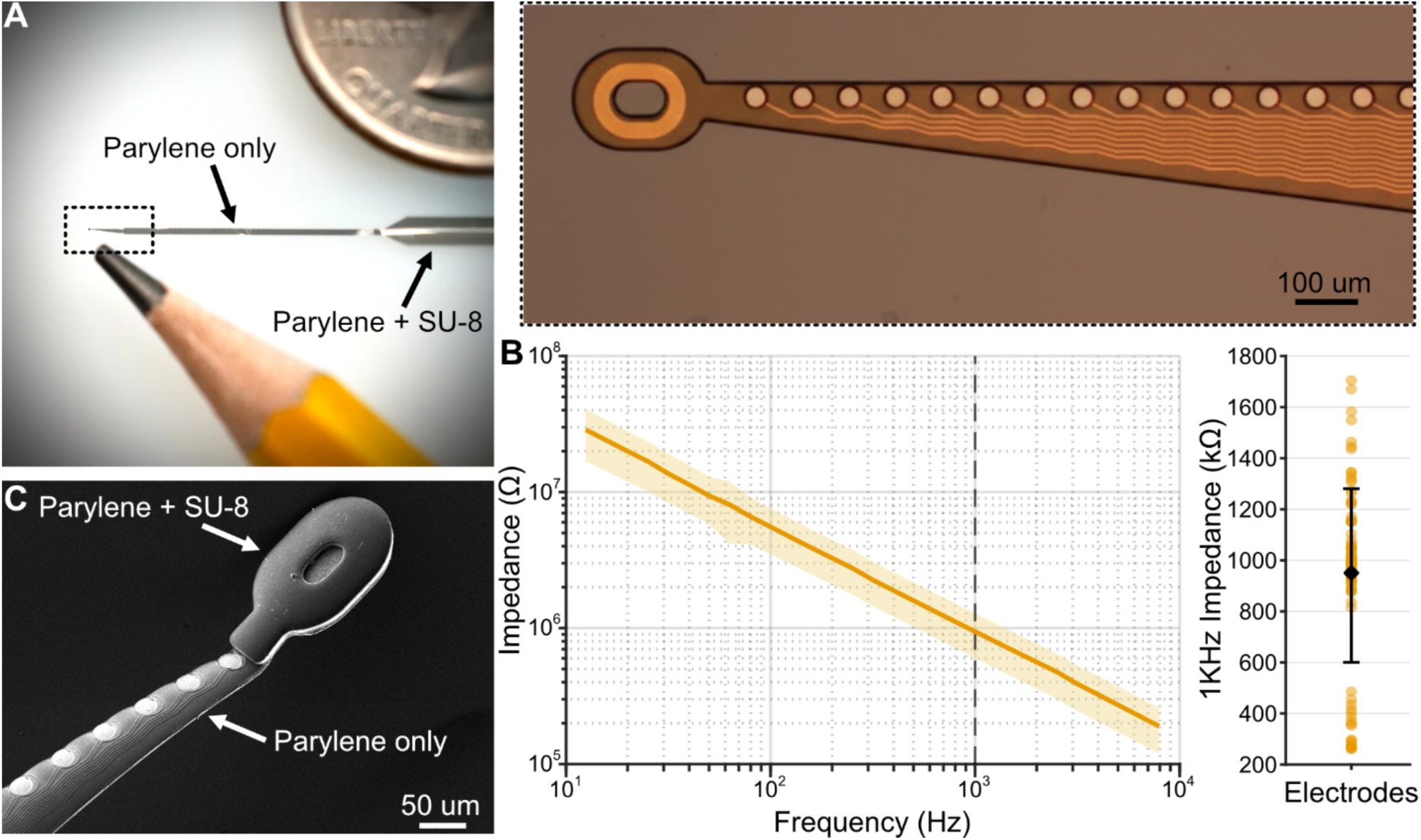
Flexible Parylene-C microelectrode arrays for NHP recordings. **(A)** Photograph of a flexible, Parylene-C probe. Inset shows microscope image of the microelectrode array. **(B)** Electrochemical impedance measurements of the Pt microelectrodes (n = 32 electrodes). Shaded area shows the standard deviation. Scatter plot shows the mean and standard deviation of the impedance at 1 KHz. **(C)** Scanning electrode micrograph of the microelectrode array and probe tip. A 10 µm SU-8 layer reinforces the 7 µm-thick Parylene-C through-hole.

Throughout the design process, we balanced the tradeoffs of ultra-flexibility with the mechanical robustness for reliable implantation in large-brained animals. The probes had an average Parylene-C thickness of 7 µm and a width of 300 µm, tapering down to 50 µm at the probe tip. To enhance mechanical stability, we added an additional 10 µm layer of SU-8 that covered the probe backend (connectors and ribbon, Fig. 1A). Similar to previous studies (Gonzales et al., 2025), we found that this dramatically reduced probe curling and eased the process of preparing probes for recordings. We also strategically placed a through-hole at the probe tip for facilitating implantation via a microwire shuttle (Luan et al., 2017; Tian et al., 2023; Y. Wang et al., 2023) and surrounded it with SU-8 to reduce the likelihood of tearing during insertion (Fig. 1C). These design features produced probes that were thin and compliant enough to move with brain tissue, yet robust enough to withstand the mechanical stresses of insertion through intact dura in large animals.

### Telescopic delivery of flexible electrodes in awake NHPs

The key technical challenges for flexible electrodes in NHPs are delivering them through thick dura and removing the stiffening shuttle. We addressed this through a two-stage “telescopic” implantation approach that reliably deployed our flexible electrodes into the awake NHP brain without the need for a new craniotomy or dura removal (Fig. 2, Methods). Our approach parallels conventional NHP neurophysiological recordings and incorporates a 19 G steel guide tube for penetration through dura/pia mater (Fig. 2A). To guide the flexible microelectrode through the guide tube and to the recording depth, our system relied on a sharpened tungsten microwire threaded through the tip of our flexible probe (Fig 2A), mirroring previous methods in rodents (Luan et al., 2017) and NHPs (Tian et al., 2023; Y. Wang et al., 2023). However, when this technique has been employed for large-brained animals, there are reports of an inability to retract the shuttle (Y. Wang et al., 2023). Often, detachment of the microwire and flexible array is unreliable, and both the shuttle and microelectrode exit tissue during retraction. To overcome this challenge, we drew inspiration from previous studies that temporarily enhance flexible electrode rigidity by supporting the flexible material as close as possible to the probe tip (X. Liu et al., 2021; Srikantharajah et al., 2021). To achieve this support during NHP insertion, we added a second “support tube” consisting of a 23 G needle (Fig. 2A). In the full system, the microwire shuttle, support tube, and guide tube fit together concentrically, creating a telescoping mechanism (Fig. 2A-B). In our design, the microwire threaded the flexible array and also formed the core of the telescope (Fig. 2A). Critically, the threaded Parylene-C probe runs along the outside of the support shuttle, pressed between the walls of the two steel tubes (Fig. 2A). Therefore, during microwire retraction, the Parylene-C was held in place by the guide tubes, enhancing rigidity, and promoting de-threading of the tungsten shuttle while the flexible array remained free-floating in brain tissue (Fig. 2B-C).

**Figure 2.**
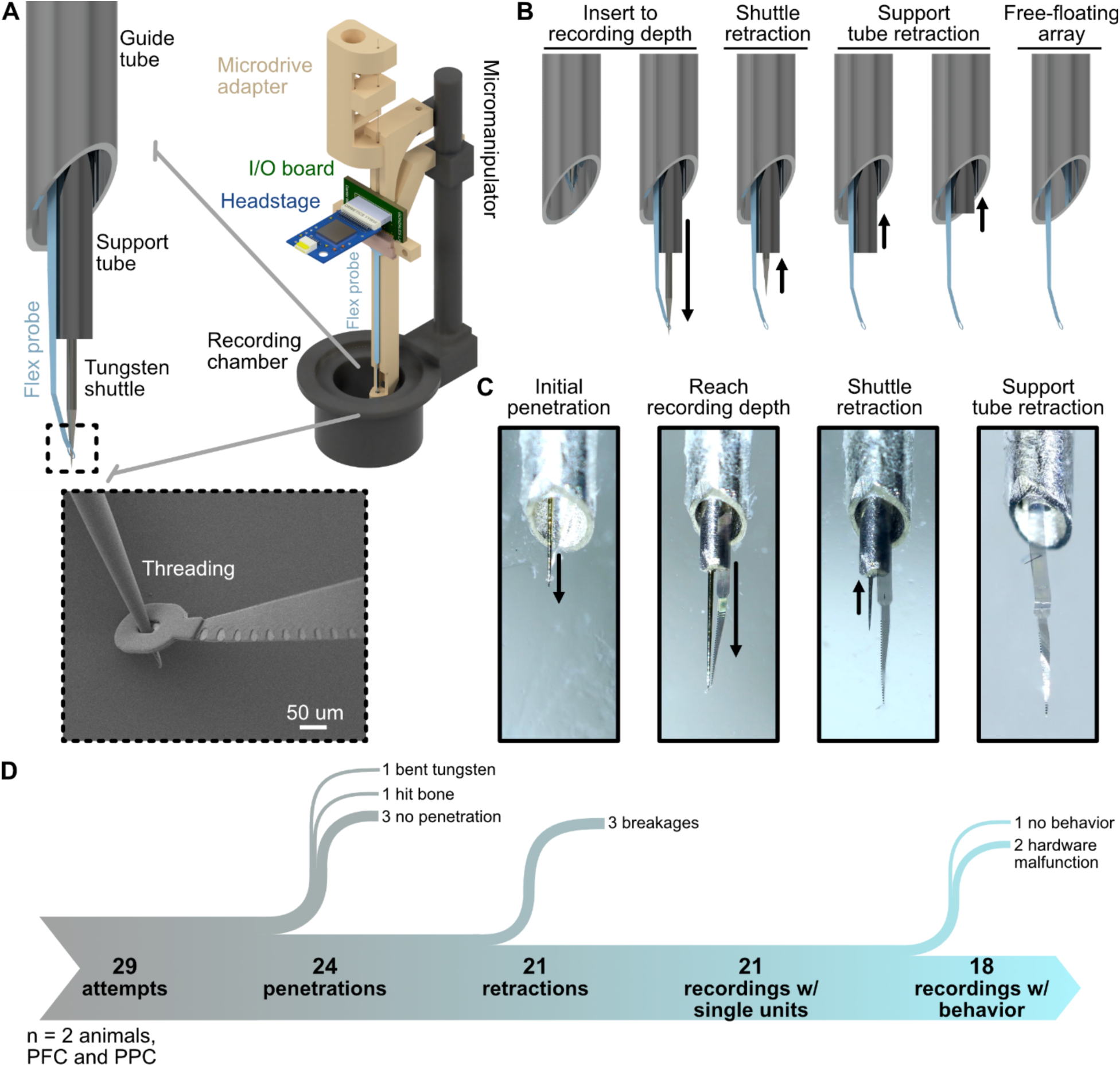
Telescopic insertion of flexible arrays. **(A)** CAD design of the full insertion system and telescoping mechanism for implanting flexible probes. Inset shows an SEM of the sharpened tungsten shuttle threading the Parylene-C through-hole. Contacts shown facing in for visualization. During recordings, probe orientation was flipped for contacts to face tissue directly. **(B)** CAD design outlining the step-by-step process of implanting flexible electrodes using telescopic insertion. **(C)** Example insertion of a flexible microelectrode array in agar. **(D)** Flow chart displaying the overall implantation outcomes and failure modes across all recording attempts (n = 2 animals).

Our full implantation assembly consisted of a commercial manipulator onto which we mounted a custom 3D-printed microdrive that housed the microelectrode array, headstage, and telescope components (Fig. 2A, Fig. S2). This yielded a total of four operable axes for independent control of each component of the telescope, enabling fine insertion of our flexible arrays and stable removal of the shuttling mechanism. Benchtop testing of this multi-step process in agar is shown in Figure 2B-C. First, we used the commercial manipulator to advance the entire system through a chronic recording chamber towards the brain until the guide tube penetrated debrided dura. Next, we used the microdrive to deliver the threaded flexible array, microwire, and support tube through the guide tube and into the brain as one unit. We inserted to a recording depth of approximately 3 mm. At this point, we retracted the tungsten microwire with a second drive block built into the adapter, leaving the flexible array and smaller support tube in the brain. Following microwire retraction, we then retracted the support tube into the larger guide tube, leaving only the flexible array in tissue at the desired recording depth (Fig 2B-C). In 29 *in vivo* attempts in two animals, we successfully penetrated to the recording depth in 24 sessions (82.8%, Fig. 2D, Table 1). Retraction was also highly reliable. Our flexible arrays remained in the brain in 87.5% of successful penetrations (21 sessions total, Fig. 2D, Table 1). In all successful retractions, we also recorded single-unit activity from the free-floating array (Fig. 2D). Critically, if any portion of the implantation failed (Fig. 2D), due to the nonsurgical setting we could simply rethread a new probe and attempt another insertion on the same or next day. These results demonstrate that our telescopic insertion process enables reliable delivery into large-brained animals through intact dura and without the need for a new craniotomy.

**Table 1.** Summary of implantation outcomes.

| **Animal ID** | **Brain Area** | **Total Attempts (N)** | **Successful Penetrations (N, %)** | **Successful Retractions (N, % of Penetrations)** | **Behavioral Recordings (N, % of Retractions)** |
| --- | --- | --- | --- | --- | --- |
| **Monkey R** | PFC | 9 | 7(77.8%) | 6(85.7%) | 5(83.3%) |
| **Monkey R** | PPC | 12 | 10(83.3%) | 9(90%) | 8(88.9%) |
| **Monkey O** | PFC | 8 | 7(87.5%) | 6(85.7%) | 5(83.3%) |
| **Overall Total** |  | 29 | 24(82.8%) | 21(87.5%) | 18(85.7%) |

### Single-unit recordings using flexible arrays in awake macaques

Having established reliable implantation procedures, we next assessed the quality and yield of single-unit activity. Our insertion process forgoes the need for new surgical interventions and therefore allowed us to perform repeated recordings in n = 2 awake, head-fixed macaques performing an oculomotor delayed response (ODR) task. We performed in vivo recordings across two brain regions, prefrontal cortex (PFC) and posterior parietal cortex (PPC), in three different recording chambers (PFC in animals R and O, PPC in animal R, Fig. 3A). Following retraction of the tungsten shuttle we stably recorded multiple neurons that exhibited robust spiking (Fig. 3B). In both brain areas, extracted waveforms and inter-spike intervals displayed single unit features (Fig. 3C-F). In total, we recorded from n = 232 putative single-units across 18 sessions (Table 2). We observed units distributed across the length of the flexible array (Fig. 3G) with an amplitude and signal-to-noise ratio (SNR) comparable to other microelectrodes (Fig. S3-4). These results suggest that the insertion process did not significantly damage tissue in the recording region.

**Figure 3.**
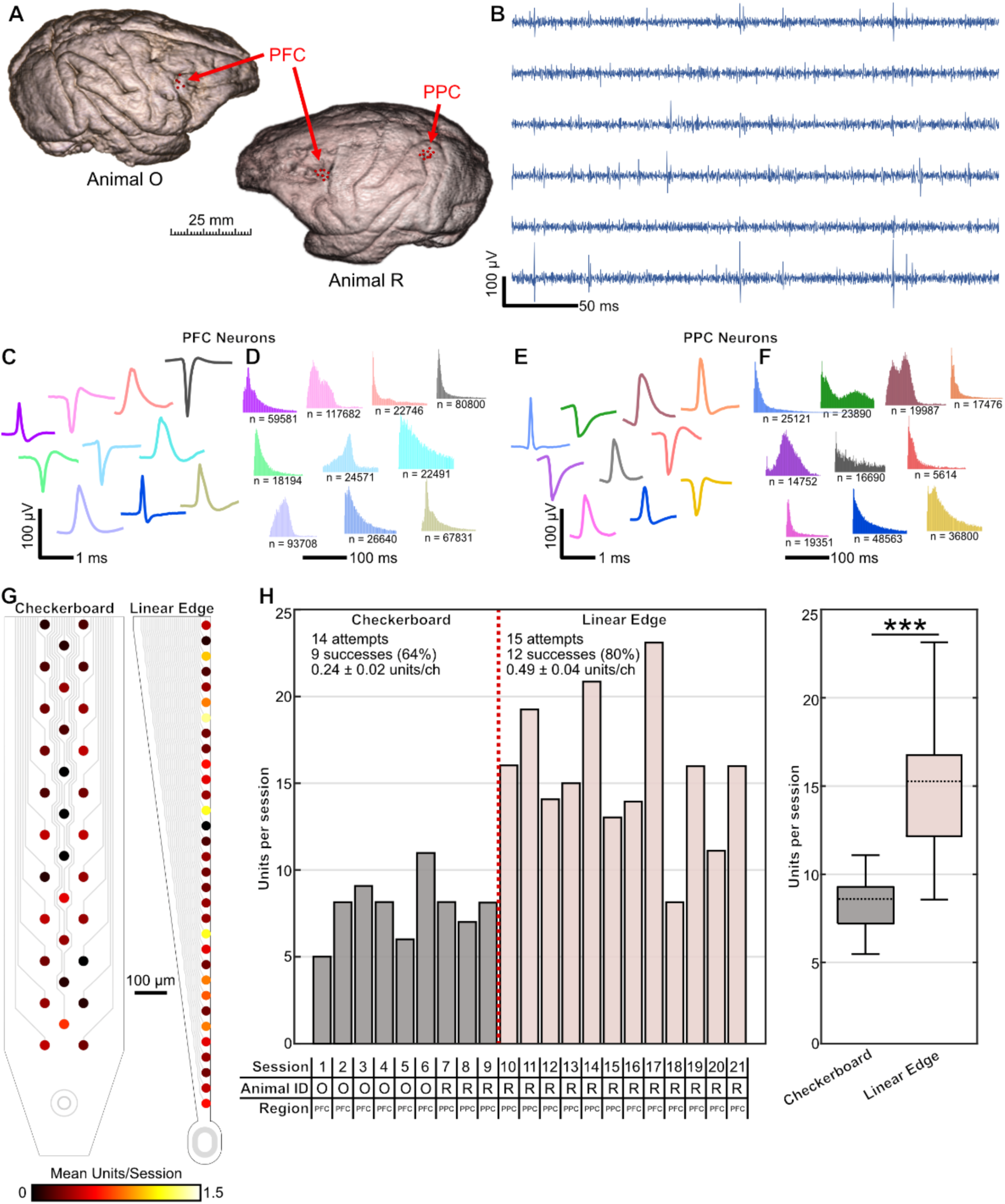
Flexible microelectrodes record single-unit activity across cortical regions. **(A)** MRI brain scans for both animals overlaid with the locations of successful recordings with flexible microelectrodes. **(B)** Representative spike band data from a select set of electrodes in the PFC of animal O. **(C-D)** Representative waveforms and inter-spike intervals detected in the PFC of animal O. **(E-F)** Representative waveforms and inter-spike intervals detected in the PPC of animal R. **(G)** Heatmap of the average number of single units detected on each recording site (n = 2 animals, 21 sessions, 232 single units). **(H)** Number of single units detected in each recording session. Sessions are organized chronologically. Red line denotes the switch from the checkerboard design to the linear edge design. Bottom table denotes the session number, animal ID, and brain area. Recording attempts, successes, and average yield per electrode are also noted. (Right) Boxplots show the distribution of units recorded per session for checkerboard and linear-edge arrays. Across sessions, linear-edge arrays yielded significantly more units per session than checkerboard arrays (15.91 ± 1.22 vs. 8.14 ± 0.59 units/session, mean ± SEM; two-sample t-test, p = 0.0002).

**Table 2.**
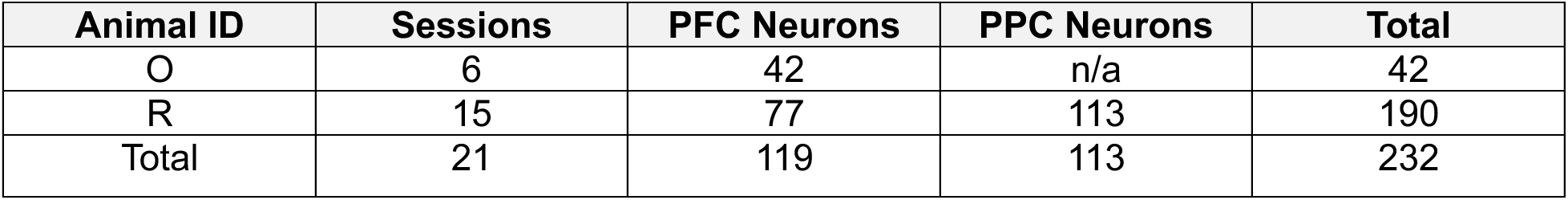
Summary of recording outcomes.

| **Animal ID** | **Sessions** | **PFC Neurons** | **PPC Neurons** | **Total** |
| --- | --- | --- | --- | --- |
| O | 6 | 42 | n/a | 42 |
| R | 15 | 77 | 113 | 190 |
| Total | 21 | 119 | 113 | 232 |

The repeatability of our implantation approach allowed us to troubleshoot different electrode array geometries for optimizing single-unit yield (Fig. 3G). We began with a three-column checkerboard design that emphasized electrode density for spike detection across multiple sites simultaneously. Surprisingly, we found that this geometry resulted in a low spike yield (Fig. 3G). We hypothesized this could be due to the electrode site distance from the probe edge, which has been observed with silicon arrays (Fiáth et al., 2021; H. C. Lee et al., 2018).

We then redesigned and fabricated a new batch of flexible arrays with a “linear edge” design, where we placed all metal contacts only 3 µm from the Parylene-C edge (Fig. 3G, Fig. 1). This geometric change immediately doubled the single-unit yield per recording site (0.24 ± 0.02 units/ch for the checkerboard design, 0.49 ± 0.04 units/ch for the linear design, mean ± SEM) and significantly increased the number of units recorded per session (Fig. 3G-H). Our recording success rate also improved throughout the study as we refined the implantation approach. Out of 12 total attempts using the linear design, we recorded single-unit activity in 9 sessions, for a success rate of 80% compared to 64% for the checkerboard experiments (Fig. 3H). Together, these results demonstrate that our telescopic insertion coupled with an optimized flexible electrode design produce high-quality recordings suitable for studying neural circuit dynamics in behaving primates.

### Flexible arrays exhibit multi-fold reductions in single-unit drift

While flexible electrodes are theorized to move with brain micromotions and reduce single-unit drift, direct quantitative comparisons with rigid probes have not been performed. Our non-surgical approach uniquely enabled us to perform this comparison by recording from the same craniotomies with both electrode types. In addition to the 21 recordings performed with flexible arrays in animals O and R, we performed 35 additional recordings in the same animals and chambers using a combination of 32 channel Plexon S/V-Probes and Diagnostic BioChips (DBC) Deep Array Probes. Both rigid probes are fabricated from stainless steel with a 200-μm shank diameter and have an electrode spacing similar to our flexible devices. For all recordings, we sorted single-units and calculated the net displacement of spikes along the probe with centroid estimation (Fig. 4A, Methods). Single-unit displacement for all rigid and flexible recording sessions is shown in Figure 4B. A dramatic reduction in drift for flexible arrays is immediately apparent. We further quantified this data by calculating the average drift velocity across 5 min bins and the maximum peak-to-trough displacement for each recording session (Fig. 4C). Both metrics indicate a multi-fold reduction in single-unit drift for flexible microelectrodes (Max displacement: 247.5, 188.9, 39.45; 5-min velocity: 67.8, 36.7, 7.45) (Fig 4C, p < 0.001, one-way ANOVA). Together, these experiments suggest that flexible arrays move with the micromotions of brain tissue and definitively prove that these state-of-the-art implants enhance recording stability compared to conventional rigid electrodes.

**Figure 4.**
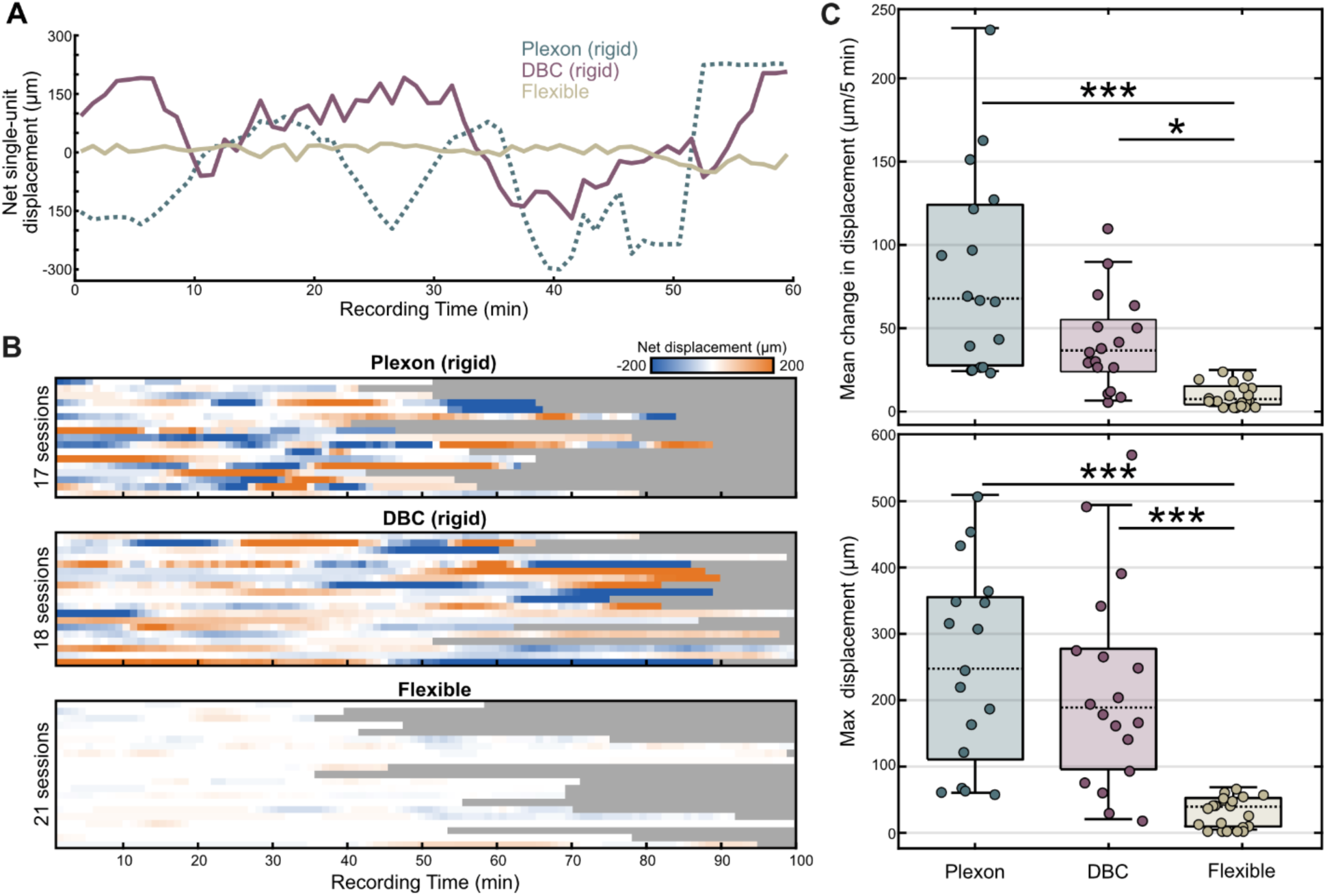
Flexible arrays have low single-unit drift compared to rigid electrodes. **(A)** Representative drift traces of 3 recordings from monkey R PPC chamber using a Plexon V probe, Diagnostic Biochips 128 channel Deep Array probe, and our flexible array. **(B)** Heatmap showing the drift activity of all Plexon, DBC, and our flexible array sessions from both monkeys O and R. Drift values are binned into 1 min windows. **(C)** (Top) Box plot summarizing the mean time derivative (i.e. drift velocity) of single-unit displacement across 5-minute windows of each session by probe type. Drift velocity for the flexible array was significantly reduced (p = 4.999 e-07, p = 0.03148 one-way ANOVA, with a post-hoc Bonferroni correction, ***p<0.0001, *p<0.05). (Bottom) Box plot summarizing maximum displacement (i.e. peak-to-trough values) of each session by probe type. Drift was significantly reduced by flexible probes (p = 1.99e-06, p = 3.321e-05, one-way ANOVA with a post-hoc Bonferroni correction, ***p<0.0001).

### Flexible arrays capture task-modulated single-unit activity during working memory

To verify that our flexible arrays capture task relevant neural activity, we analyzed single-unit responses during the ODR task (Fig. 5A). The ODR task is a well-established approach for probing working memory, which is a significant function of PFC and PPC (Constantinidis et al., 2018; Katsuki and Constantinidis, 2012). From 21 neural recording sessions, 18 also contained sufficient behavioral data during execution of this task (Fig. 2D). We classified single units into four commonly identified subgroups: cue responsive, delay responsive, saccade responsive, or responsive to multiple task epochs within the same trial, based on their firing rate (see Methods). Furthermore, we identified neurons showing tuning to location during the delay epoch, as PFC neurons are known to do (Constantinidis & Goldman-Rakic, 2002).

**Figure 5.**
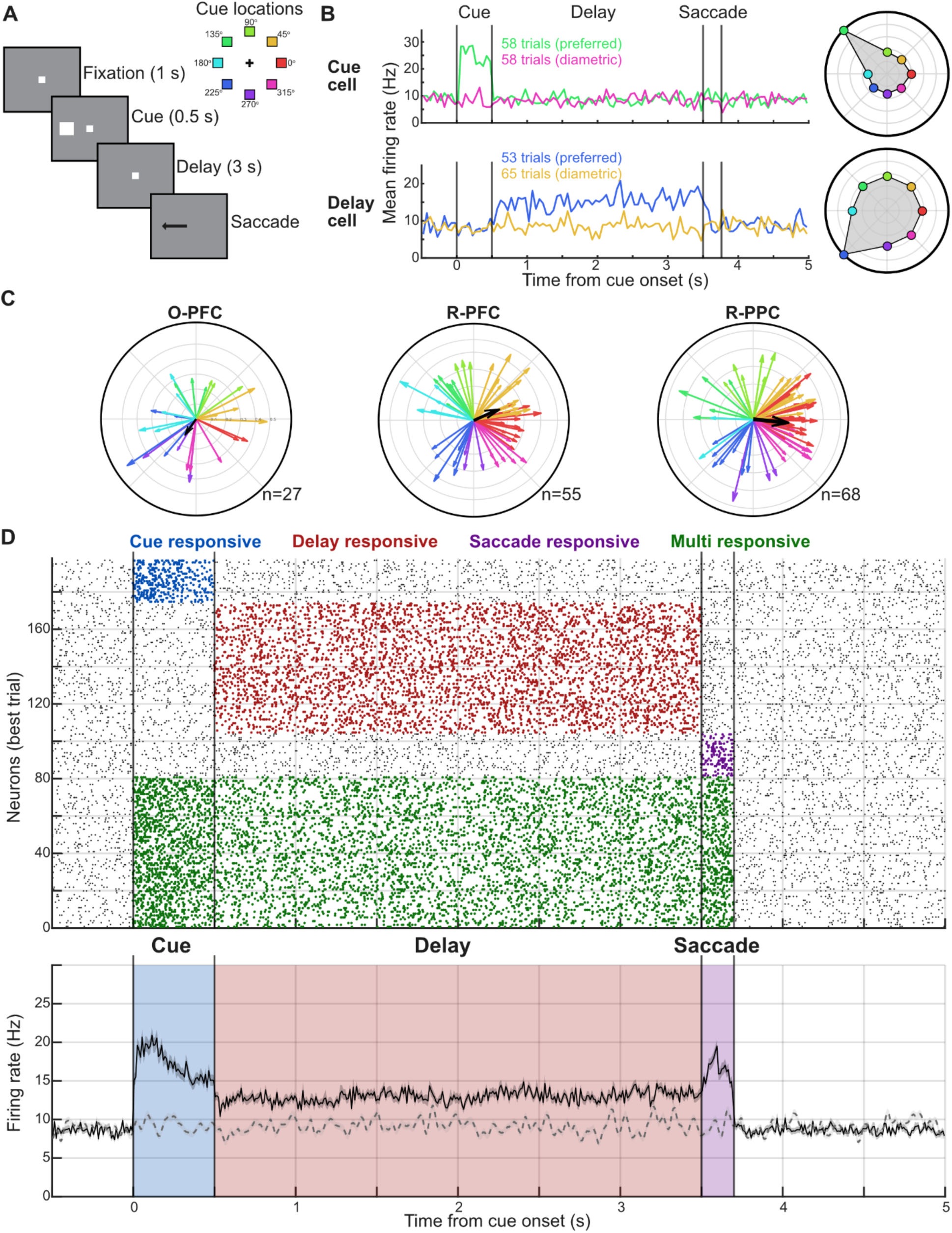
Task relevant activity recorded from flexible array. **(A)** Left: Sequence of events in the ODR task. The monkey is required to maintain fixation for 1 s to observe the cue stimulus, remain fixated for the delay period of 3 s, and make a saccade to the remembered cue location after the delay period, indicated by the disappearance of the fixation point. The monkeys were rewarded for the successful completion of the trial. Right: Cue stimuli locations are shown in a ring with different colors. **(B)** Representative cue-response (top) and delay-responsive (bottom) cells. Plots show the peristimulus time histogram (PSTH) for trials with the cue in the preferred location compared to the target 180° away in the diametric location. Compass plots show cell tuning. **(C)** Compass plot showing tuning of all single units selective for the delay period recorded in each brain area. Each arrow’s direction denotes the angle of tuning and length indicates the strength of tuning for each unit. Each arrow color indicates the spatial selectivity of that neuron. **(D)** Top: Raster plot showing single trial activity of all single units that showed behaviorally relevant firing, organized as cue, delay, saccade, and multiple epoch responsive units. Bottom: PSTH for the trials shown in the raster above, across all single units. Solid trace shows the average activity for all units’ preferred cue location. Dashed line shows average activity for all units for the diametric location.

From 232 total single units recorded across sessions in PFC and PPC, 197 were identified as task-responsive (see Methods). These included 23 cue-responsive, 70 delay-responsive, 23 saccade-responsive, and 81 multi-epoch neurons. We quantified the spatial tuning of these cells during the delay period and found strong single-neuron tuning distributed across target locations (Fig. 5B-C), with a slight bias in population tuning towards the contralateral field-of-view (Fig. 5C). The distribution of functional cell types did not differ between PFC and PPC (χ²(4) = 1.12, p = 0.89). We compiled a population-averaged firing rate for all task-responsive cells (Fig. 5D). Individual neurons maintained consistent firing patterns across the recording session (Figure S5-6). These results confirm that our flexible arrays reliably capture task-modulated neural activity with response properties characteristic of PFC and PPC, demonstrating their suitability for studying cognitive neuroscience in behaving primates.

## DISCUSSION

We successfully developed a flexible electrode array design suitable for NHP recordings and demonstrated a non-surgical, repeatable approach for inserting them in awake, behaving NHPs. We attribute our improved success over previous studies to our telescopic approach for implantation, which reliably delivered free-floating flexible arrays through intact dura. This insertion approach allowed us to optimize the electrode array design to maximize single-unit yield. In addition, for the first time, we quantitatively showed that highly flexible electrode arrays have superior single-unit recording stability compared to traditional rigid electrodes. Recording stability improves the fidelity of single-neuron firing rate measures, such as those we quantified in our study, but also measures of synchronous activity between neurons, such as spike-count correlation and cross-correlation (Qi & Constantinidis, 2012). Combined, our methods led to weeks of repeated recordings across multiple brain regions in NHPs performing cognitive tasks and hundreds of neurons associated with working memory.

Our work substantially lowers barriers to flexible electrode adoption in the broader NHP neurophysiology community. Our non-surgical, acute approach provides an accessible entry point for laboratories to adopt flexible electrode technology without high-risk surgeries. Our approach works through standard recording chambers with intact dura, allowing researchers to perform flexible electrode recordings using the same infrastructure and workflows they already use for rigid probes. We used a combination of a commercial micromanipulator, off-the-shelf parts, and a 3D printed microdrive with designs that we have made freely available (Methods). This compatibility means laboratories can integrate flexible arrays without major equipment investments, specialized surgical training, or procedural overhauls. In addition, this also means that animals with recording chambers that are not currently being used for active studies can be repurposed for flexible electrode testing. In addition to the practicality of our approach, previous flexible electrode demonstrations in NHPs have not reported the outcomes of all implantation attempts, suggesting that recordings were not only high-risk but potentially unreliable. To the best of our knowledge, our study is the first to report all implantation outcomes (Fig. 2D). From 29 total attempts in awake animals, we achieved single-unit recordings in 21 sessions (72% success). Notably, after optimizing the probe design, we successfully implanted and recording in 12 out of 15 attempts (80% success, Fig. 3H), and we expect this yield to increase as we continue to minimize human error (see Supplementary Discussion). These data provide NHP groups clear expectations for success rates and failure modes, enabling realistic planning and iterative improvement rather than reliance on selectively reported successes. Finally, we showed the first side-by-side recording comparisons with traditional rigid microelectrode arrays. Reduced scarring and gliosis from polymer electrodes has been extensively reported by previous studies (Chung, Joo, Fan, et al., 2019; Kim et al., 2013; Lu et al., 2016; Luan et al., 2017; Musk & Neuralink, 2019; Yasar et al., 2024; S. Zhao et al., 2023; Z. Zhao et al., 2022; Zhou et al., 2017). Here, we show that free-floating flexible arrays also have superior single-unit tracking stability, likely because these tissue-like interfaces move with natural brain micromotion. Combined, our work provides the technical innovation to overcome implantation challenges and the quantitative evidence to validate theoretical advantages, further establishing flexible microelectrode arrays as a practical, accessible technology for NHP neuroscience.

Importantly, our acute insertion method establishes the foundation for a developing a platform for chronic, large-scale, and highly stable longitudinal tracking of single neurons in large brains. While the flexible arrays we used in this study have only 32 recording sites, the microfabrication process is scalable and can be increased to hundreds of channels with no additional cost or time (Methods). Our results suggest that scaling comes with important design considerations, such as metal contact placement and overall shank width. Increasing the electrode count also non-trivially increases the complexity and size of the I/O interface, which should be kept small to easily anchor to the recording chamber. With these considerations, we believe scaling to a double-sided,128-channel shank is feasible and that implanting multiple of these arrays simultaneously can be achieved for regular use in an academic NHP laboratory.

For troubleshooting chronic implants, our platform has unique advantages compared to relying on a new craniotomy. For example, animals with existing craniotomies can be used for developing chronic techniques, rather than risking a new craniotomy purely for testing long-term implantation. In addition, techniques for anchoring electrodes in tissue can be tested for days or weeks at a time, followed by removal and re-implantation of a new array.

Overall, our study demonstrates that flexible electrodes can be reliably implanted with high yield in behaving NHPs and provide substantially improved short-term stability relative to rigid microelectrodes. Although our experiments are acute, this level of stability is a critical prerequisite for chronic implementations, establishing a practical path toward tracking large neuronal populations across extended timescales. By lowering the technical barriers to reliable flexible-array recordings, our approach lays the groundwork for future studies that can follow how individual neurons and population-level computations evolve over learning, development, and cognitive change.

## METHODS

### Flexible Microelectrode Fabrication

We developed a custom design for fabricating ten Parylene-C–based probes simultaneously on 4-inch silicon (Si) wafers using a standard photolithography process. We first cleaned each wafer by soaking it for 5 minutes in acetone, followed by 5 minutes in isopropyl alcohol (IPA). Afterwards, we dried the wafers with nitrogen and then O₂-plasma cleaned them for 1 minute (Trion – Phantom RIE). We deposited ∼3 µm of Parylene-C using chemical vapor deposition (Specialty Coating Systems) with 3.5 g of dimer for four wafers at once. To improve surface adhesion and remove any residual monomers, we vacuum-baked the samples overnight at 160 °C at 30 mm Hg.

We used mask photolithography to pattern metal recording contacts, leads, connection pads, and the through-hole at the probe tip. We repeated the cleaning sequence (IPA, acetone, drying, and 10 second O₂ plasma) to prepare the Parylene-C surface. A 450 nm layer of LOR7A lift-off resist was spin-coated and baked at 160 °C for 10 minutes. Subsequently, a 1.3 µm layer of S1813 was spin-coated and baked at 115 °C for 1 minute. Using a mask aligner (Karl Suss Ma-6), we exposed the photoresist with the metal pattern at 110 mJ/cm² and developed it for 1 minute in MF-319 (Kayaku), then rinsed it in deionized (DI) water. We O₂-plasma cleaned the wafers for 10 seconds to improve metal adhesion, then deposited 200 nm of Pt at ∼2.5 nm/min using DC sputtering (AJA ATC-2200). We lifted off the metal layer at 80 °C then at room temperature overnight in Remover 1165 (Microposit), then rinsed in DI water. We O₂-plasma cleaned the surface again for 10 seconds to remove any solvent residue and reactivate the surface.

We encapsulated the patterned metal layer by depositing another ∼3 µm of Parylene-C using the same method, then vacuum-baked the wafers again overnight at 160 °C and 30 mm Hg. To define the recording sites, connection pads, and probe outline, we first deposited a photoresist etch mask onto the parylene-Pt-parylene stack. We began by repeating the cleaning steps, then spun 12 µm of SPR 7.0-220 (Kayaku) and prebaked the sample at 70 °C for 150 seconds, then at 115 °C for another 150 seconds. After cooling for 3 minutes, we exposed the resist twice at 280 mJ/cm²—once with a mask revealing the contacts and pads, and once for the probe outline. We let the sample rest for 35 minutes, post-baked it at 70 °C for 90 seconds, then at 115 °C for 90 seconds, and rested it again for 10 minutes. We developed the resist in MF-319 for 5 minutes.

We etched the Parylene-C using high-power O₂ plasma (250 W RIE, Trion Minilock) to expose the metal contacts (below the top 3 µm layer of parylene) and define the probe outline (through 6 µm of parylene) simultaneously. To prevent overheating, we alternated 2 minutes of etching with 1 minute of cooling, repeating this cycle 10 times for a total etch time of 20 minutes, or until the Si wafer was visibly clear of Parylene-C. To avoid over-etching, we designed the recording-site etch mask with a 25 µm diameter—5 µm smaller than the metal layer—which expanded to a final diameter of 30 µm during etching. Afterward, we stripped the remaining photoresist in acetone, rinsed in IPA, and dried the wafers with nitrogen.

To improve handling of the long, thin polymer devices, so we added a photolithography step to thicken the probe backend. We activated the Parylene-C surface with a 30-second O₂-plasma clean, spun SU-8 2010 (Kayaku) to ∼12 µm thickness, and baked the wafers for 3 minutes at 95 °C. Using a mask aligner, we exposed the resist with 135 mJ/cm², baked for 4 minutes at 95 °C, developed in SU-8 developer for 3 minutes, and rinsed in IPA. To remove microcracks, reduce stress, and improve adhesion to Parylene-C, we hard-baked the wafers in a stepped ramp: 2 minutes each at 100 °C, 110 °C, and 120 °C, followed by 1 hour at 130 °C. This SU-8 layer reinforced the backend and surrounded the probe’s through-hole tip, while the implantable shank remained Parylene-C only, preserving flexibility.

### Probe Packaging

We released the Parylene-C probes by submerging the wafer in DI water and peeling the devices using fine-tip surgical tweezers. To minimize damage to the conducting layer, we designed a small handle the backend region that included only SU-8 and Parylene-C with no platinum, and we used this area to begin peeling. Once the handle separated from the wafer surface, the rest of the device released easily from the wafer. The SU-8 layer reduced curling compared to 6 µm-thick Parylene-C probes and made handling easier. In an effort to minimize curling further, we annealed the probes between Teflon sheets using a vacuum bake at 160 °C for 2 hours, at a pressure of 30mmHg (Thielen & Meng, 2023).

After drying, we bonded the probe backend to a laser-cut 0.010-inch-thick polyether-ether-ketone (PEEK) film (CS Hyde, 37-10F-24” × 24”) using superglue, and any excess Parylene-C was trimmed off the backend (Gutierrez et al., 2011). This PEEK support allowed stable insertion and clipping into a 32-channel zero-insertion-force (ZIF) connector (Hirose FH12-32S-0.5SH) mounted on a custom 1.0 × 0.5-inch printed circuit board (PCB). We soldered a ground pin and a 32-channel Omnetics connector (A79022-001) to the PCB, allowing it to interface with a 32-channel Intan amplifier (RHD 2132).

### Electrochemical Impedance

We tested impedance using three-electrode electrochemical impedance spectroscopy in PBS saline (Gamry Interface 1010E, 10 mV RMS amplitude, 1-10,000 Hz sweep). The recording site was used as the working electrode, an Ag/AgCl electrode as the reference electrode, and Pt wire as the counter electrode. We also had the ability to test the 1 KHz impedance using on-chip measurements with the Intan headstage.

### Experimental Model and Subject Details

Two male, ∼9 years old rhesus monkeys (*Macaca mulatta*) weighing 9-14 kg were used in this study. Monkeys were single housed in communal rooms, and they had sensory interactions with other monkeys. All surgical and animal use procedures were reviewed and approved by the Vanderbilt University Institutional Animal Care and Use of Laboratory Animals and the National Research Council’s Guide for the Care and Use of Laboratory Animals.

### Surgery

Animals were implanted with a 20-mm-diameter recording cylinder during previous studies (Zhu et al., 2023, 2025), and we continued the use of these chambers for the current study. Both animals used for recordings in this study had a chamber over the dorsolateral prefrontal cortex (Animals R and O) and one subject had a posterior parietal cortex chamber (Animal R). We registered CT and MR images to localize the cylinder and electrode penetrations. We aligned anatomical T1 images to the high-resolution NIH Macaque Template (NMT) using @animal_warper (Jung et al., 2021; Saad et al., 2009). We used the 3D slicer software to find the projection of the recording cylinder on the brain surface, then localized electrode penetrations using the grid locations relative to the recording cylinder.

PFC for both animals targeted the dorsolateral region (Broadmann areas 8a and 46) along the bank of the principal sulcus using preregistered MRI scans with chamber coordinates mapped to the brain. PPC recordings primarily targeted the intraparietal sulcus using the same registration with chamber coordinates.

### Flexible Electrode Implantation

Advancing our flexible microelectrodes into the brain required a custom “microdrive” adapter that we attached to a conventional micromanipulator (MO-97A, Narshige Corp.). We designed and 3D printed the microdrive similar to other instrument designs (Bauer et al., 2023; Q. Wang et al., 2021). The designs are openly available online (https://github.com/GonzalesLabVU/NHPflex2025).

The Narishige microdrive consists of a dovetail interface between the two rails of our 3D printed adapter: a main driver that held a sharpened 19 G guide tube and a second driver that that mounted our flexible probe, tungsten wire, support tube, PCB, and headstage. Each rail could be operated independently. We used the main driver to lower the entire assembly towards the brain for initial penetration of the dura using the guide tube. We followed this by using the second rail for fine scale driving of the flexible probe threaded to a tungsten microwire (FHC Inc., UEWLGGSE4N1E) and encased by the 23 G blunt support tube. After advancing the probe tip to the desired recording depth (∼3000 µm), we used our third custom driver that retracted the tungsten microwire from the brain via a screw. The 23 G encasement facilitated detachment of the tungsten and Parylene-C. Once the tungsten microwire was removed, we retracted the 23 G encasement tube into the 19 G guide tube, leaving the flexible microelectrode freely in cortical tissue. The guide tube remained inserted in dura for the duration of the recordings Threading of the flexible array occur prior to implantation. We first pre-routed the tungsten microwire through microdrive and 23 G support tube, then threaded the microwire tip to the flexible array and temporarily attached the two with the assistance of poly-ethylene glycol (PEG). We aligned the microwire such that it protruded from the support tube by approximately 5 mm.

### Neurophysiological Recordings

We collected electrophysiological recordings using our custom flexible microelectrodes with an Intan RHD 2132 headstage. For rigid recordings, we used both Plexon S/V probes and Diagnostics BioChips 128 Deep Array probes. The Plexon probes used Intan RHD 2132 headstages, while the DBC probes have a custom headstage that utilizes Intan RHD amplifiers. As with the flexible arrays, all rigid electrodes were implanted using a conventional micromanipulator (MO-97A, Narshige Corp.) and advanced through a guide tube. All recordings used an OpenEphys data acquisition system (OpenEphys) sampled at 30 KHz.

### Behavioral Task

The monkeys sat head-fixed in a primate chair while viewing a 1920 × 1080, 60 Hz monitor (Samsung QMC32C) positioned 69 cm from their eyes under dim ambient light. An infrared eye tracker (ISCAN ETL-200; ISCAN, Inc.) monitored eye position at 500 Hz with at least 0.5° precision at the center point. We implemented visual stimulus control, eye monitoring, and synchronization with neural data in MATLAB (MathWorks) using the Psychophysics Toolbox (Meyer & Constantinidis, 2005).

Animals were previously trained on an oculomotor delayed response (ODR) task (Funahashi et al., 1989). Each trial began with fixation on a 0.1° white square at the screen center. We required fixation within a 3° window around the central point while presenting peripheral visual stimuli at radial locations. If fixation broke at any time, we terminated the trial and withheld reward. For the cues, a 1° white square appeared for 0.5 s at one of eight locations spaced by 45° around a 10° eccentric circle. The monkeys had to remember the cue location during a 3 s delay period. When the fixation point disappeared, they had 0.6 s to make a saccade to the remembered location and then hold fixation for 0.1 s within a 6° radius window centered on the cue. Correct trials earned a water reward. Animals typically performed between 400-500 trials over a 1-hour recording session, where they achieved a performance of approximately 90% correct trials.

## QUANTIFICATION AND STATISTICAL ANALYSIS

### Neural Data Processing

All neurophysiological analysis and quantification were performed using MATLAB. These sets of analyses use the same methods as our previous studies on working memory (Mozumder et al., 2024, 2026; Mozumder & Constantinidis, 2023; Thrower et al., 2023; Z. Wang et al., 2022; Zhu et al., 2023). All data was first bandpass filtered from 500-8000 Hz. We identified spike waveforms and sorted single units using the automated spike-sorting software Kilosort 2.5 (rigid electrodes) and Kilosort 4 (flexible electrodes) (Pachitariu et al., 2024) (https://github.com/MouseLand/Kilosort) and manually curated the clustered data using Phy2 (https://github.com/cortex-lab/phy/). These Kilosort versions use the same drift calculations (Pachitariu et al., 2024).

### Spike Signal-to-Noise Ratio (SNR) Analysis

Spike signal-to-noise ratio (SNR, Fig. S3) was quantified based on the ratio of spike amplitude to background noise (Jun et al., 2017). For each unit, SNR was defined as:

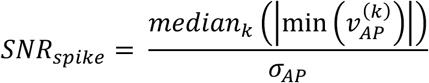

Where:

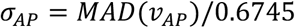

And:

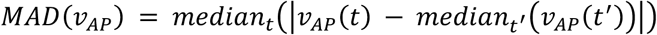

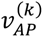 is the band-pass filtered (300-8000 Hz) action-potential waveform for spike *k*, aligned to the spike time. The median across spikes was used to provide robustness to outliers. 𝜎_AP_is the background noise and estimated from randomly selected, spike-free segments of the recordings. *MAD* denotes the median absolute deviation computed from background activity.

### Probe Drift Calculation

Drift (Fig. 4) was quantified directly from sorted spike data exported from Kilosort for each recording session using the three probe types (our flexible arrays, Plexon S/V-probes, and Diagnostic Biochips Deep Array 128 channel probe). These values have been reliably used for drift comparisons in several previous studies (Trautmann et al., 2025). The analysis was performed using custom scripts in MATLAB (MathWorks). For each probe during a recording, a drift trace was constructed to represent the temporal displacement of the population spike depth centroid over time (Steinmetz et al., 2019), expressed in µm. Template depth was determined as the signal power-weighted center of mass (COM) of the template waveforms across recording channels (Pachitariu et al., 2016). For each template *k*, the signal power on channel *c* was computed as the sum of squared waveform amplitudes, and the corresponding depth was obtained as 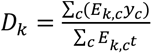 where *E_k,c_* is the channel’s vertical position. Each spike was then assigned a depth 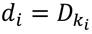 according to its template identity. Spike times were divided into consecutive temporal bins of 60 s, and for each bin *b* containing *N*_b_ spikes (minimum 25 spikes/min), the median spike depth was computed as the bin-wise center of mass: *COM*_1_ = *median*{*d_i_*: *i* ∈ *B_b_*}. The resulting time series depth centroids *COM*(*t_b_*) reflects the population’s median depth as a function of time. Net drift displacement was defined as the total range of motion across the session *D_net_* = max(*COM*) − min (*COM*). This metric quantifies overall amplitude of the probe-relative movement *D_net_*.

To quantify the temporal change of probe motion, we computed a drift “velocity” metric. Using the same median-centered spike depth centroid trace *COM*(*t*) described above, drift velocity was defined as the mean absolute temporal derivative of the probe position evaluated at a 5-min resolution. Specifically, the centroid trace was sampled every five consecutive 1-min bins yielding a down sampled sequence *COM*_5_(*k*). Drift velocity was then computed for each recording as:

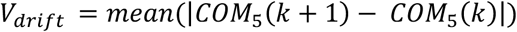

To visualize probe drift dynamics across all recording sessions, a heatmap representation of median-center spike depth drift was constructed as follows. For each session, the median centered spike depth drift trace was summarized by 1-minute non-overlapping bins. Within each bin the median drift value across all samples within was computed. All sessions were capped at 100 minutes for session wise drift analysis. The box plots summarizing distribution of drift velocity and peak-to-trough median centered spike drift trace per session were aggregated, drawing interquartile range and whiskers extending to 1.5x the IQR.

### Spike Amplitude and Single-Unit Yield

For each unit, we used the spike times from Kilosort to extract a 1 ms window around each detected spike. We bandpass filtered these segments 500-8000 Hz and common average referenced to reduce any correlated noise. Spike amplitude was calculated as peak to trough voltage displacement during the window for each unit’s spike. Unit yield was quantified at both the channel and session levels. Channel-level yield was defined as the number of units assigned to each recording site. Session yield was defined as the total number of units detected during a recording session.

### Neural Firing Analysis

We generated single-trial peristimulus time histograms (PSTHs) for visualizations by calculating spiking events of single units in 50 ms bins. We defined neurons as selective during any task epoch if they showed significantly different responses to the spatial location of the stimulus via a one-way ANOVA test on the firing rates across trials during that specific task epoch. In order to avoid false positives with sparsely spiking cells, we required that a selective neuron exhibit a firing rate of at least 4 spikes per second for its best stimulus location during the task period where the ANOVA test indicated a significant main effect.

To identify each neuron’s preferred location, we computed the circular mean of the cue angles weighted by the neuron’s mean spike count during the delay period (Wimmer et al., 2014). For each neuron, we computed:

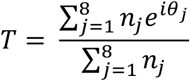

Where 𝑛_8_ is the mean spike count during the delay period in response to the cue 𝜃*_j_* (𝑗 = 1 … 8) and we extracted its modulus *T* and angle 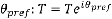. The angle 𝜃*_pref_* constitutes our estimate of the neuron’s preferred location during the delay.

### Population Firing Rate

To illustrate how the population firing rate evolved over time (Fig. 5), we averaged the activity of all neurons recorded from all sessions for each neuron’s preferred cue location (the preferred cue for each neuron was chosen as mentioned in the above section). In other words, we aligned each neuron’s activity across all the trials for the preferred location which yielded the mean population firing rate (Tang et al., 2019).

## STATISTICAL ANALYSIS

All statistical testing was performed in MATLAB. For boxplots, the upper and lower edges of the box represent the upper and lower quartiles of the data, respectively, and the line within the box represents the median. Whiskers extend to the most extreme data points not considered outliers; points beyond the whiskers are plotted individually as outliers.

Data were tested for normality using a Kolmogorov–Smirnov test. The presented data did not violate assumptions of normality so parametric statistical tests were used. Specifically, two-sample t-tests were used to compare unpaired data between two groups, and results are reported as mean ± SEM unless otherwise noted. Effect sizes for pairwise comparisons were quantified using Cohen’s d. We used a one-way ANOVA to compare unpaired data between more than 2 groups and followed this with a post-hoc Bonferroni test to correct for multiple comparisons.

## DATA AVAILABILITY

Data, code, probe layout files, and microdrive design files will be made publicly available upon acceptance.

## ACKNOWLEDGEMENTS

This work was funded by the Howard Hughes Medical Institute Hanna Gray Fellowship GT16792 (DLG), Burroughs Wellcome Fund G-1021695.01 (DLG), R01 EY017077 (CC), and R01 EY036089 (CC). All devices were fabricated and characterized using facilities within the Vanderbilt Institute for Nanoscale Science and Engineering (VINSE). We thank the VINSE staff for their infrastructure and training support in addition to Owen Meilander for assistance with sample fabrication. We thank Jaela Bills and Kayla Yetman for technical help with experiments.

## AUTHOR CONTRIBUTIONS

DG and CC designed the experiments. DW and GA fabricated the device. DW, RM, WD, and AC performed behavioral training and neurophysiological recordings. DW and RM performed data analyses. DW and DG wrote the paper with input from all authors. DG and CC oversaw the work.

## SUPPLEMENTAL INFORMATION

Document S1: Supplementary Discussion. Figures S1-S6.

## REFERENCES

Bastos, A. M., Loonis, R., Kornblith, S., Lundqvist, M., & Miller, E. K. (2018). Laminar recordings in frontal cortex suggest distinct layers for maintenance and control of working memory. Proceedings of the National Academy of Sciences, 115(5), 1117–1122. 10.1073/pnas.1710323115

Bauer, D. L., Pobiel, B., Hilber, K., Verma, A. K., Wang, J., Vitek, J., Johnson, M., & Johnson, L. (2023). 3D printed guide tube system for acute Neuropixels probe recordings in non-human primates. Journal of Neural Engineering, 20(3), 036009. 10.1088/1741-2552/acd0d7

Boussard, J., Windolf, C., Hurwitz, C., Lee, H. D., Yu, H., Winter, O., & Paninski, L. (2023). DARTsort: A modular drift tracking spike sorter for high-density multi-electrode probes. bioRxiv, 2023.08.11.553023. 10.1101/2023.08.11.553023

Carmena, J. M., Lebedev, M. A., Crist, R. E., O’Doherty, J. E., Santucci, D. M., Dimitrov, D. F., Patil, P. G., Henriquez, C. S., & Nicolelis, M. A. L. (2003). Learning to Control a Brain–Machine Interface for Reaching and Grasping by Primates. PLoS Biology, 1(2), e42. 10.1371/journal.pbio.0000042

Chung, J. E., Joo, H. R., Fan, J. L., Liu, D. F., Barnett, A. H., Chen, S., Geaghan-Breiner, C., Karlsson, M. P., Karlsson, M., Lee, K. Y., Liang, H., Magland, J. F., Pebbles, J. A., Tooker, A. C., Greengard, L. F., Tolosa, V. M., & Frank, L. M. (2019). High-Density, Long-Lasting, and Multi-region Electrophysiological Recordings Using Polymer Electrode Arrays. Neuron, 101(1), 21–31.e5. 10.1016/j.neuron.2018.11.002

Chung, J. E., Joo, H. R., Smyth, C. N., Fan, J. L., Geaghan-Breiner, C., Liang, H., Liu, D. F., Roumis, D., Chen, S., Lee, K. Y., Pebbles, J. A., Tooker, A. C., Tolosa, V. M., & Frank, L. M. (2019). Chronic Implantation of Multiple Flexible Polymer Electrode Arrays. Journal of Visualized Experiments : JoVE, 152, 10.3791/59957. https://doi.org/10.3791/59957

Chung, J. E., Sellers, K. K., Leonard, M. K., Gwilliams, L., Xu, D., Dougherty, M. E., Kharazia, V., Metzger, S. L., Welkenhuysen, M., Dutta, B., & Chang, E. F. (2022). High-density single-unit human cortical recordings using the Neuropixels probe. Neuron, 110(15), 2409–2421.e3. 10.1016/j.neuron.2022.05.007

Constantinidis, C., Funahashi, S., Lee, D., Murray, J. D., Qi, X.-L., Wang, M., & Arnsten, A. F. T. (2018). Persistent Spiking Activity Underlies Working Memory. The Journal of Neuroscience, 38(32), 7020–7028. 10.1523/JNEUROSCI.2486-17.2018

Constantinidis, C., & Goldman-Rakic, P. S. (2002). Correlated Discharges Among Putative Pyramidal Neurons and Interneurons in the Primate Prefrontal Cortex. Journal of Neurophysiology, 88(6), 3487–3497. 10.1152/jn.00188.2002

Fiáth, R., Meszéna, D., Somogyvári, Z., Boda, M., Barthó, P., Ruther, P., & Ulbert, I. (2021). Recording site placement on planar silicon-based probes affects signal quality in acute neuronal recordings. Scientific Reports, 11(1), 2028. 10.1038/s41598-021-81127-5

Forrest, A. M., Kunigk, N. G., Collinger, J. L., Gaunt, R. A., Chen, X., Vande Geest, J. P., & Kozai, T. D. Y. (2025). Finite element model predicts micromotion-induced strain profiles that correlate with the functional performance of Utah arrays in humans and non-human primates. Journal of Neural Engineering, 22(6), 066008. 10.1088/1741-2552/ae1bda

Funahashi, S., Bruce, C. J., & Goldman-Rakic, P. S. (1989). Mnemonic coding of visual space in the monkey’s dorsolateral prefrontal cortex. Journal of Neurophysiology, 61(2), 331–349. 10.1152/jn.1989.61.2.331

Gerbella, M., Borra, E., Pothof, F., Lanzilotto, M., Livi, A., Fogassi, L., Paul, O., Orban, G. A., Ruther, P., & Bonini, L. (2021). Histological assessment of a chronically implanted cylindrically-shaped, polymer-based neural probe in the monkey. Journal of Neural Engineering, 18(2), 024001. 10.1088/1741-2552/abdd11

Gilletti, A., & Muthuswamy, J. (2006). Brain micromotion around implants in the rodent somatosensory cortex. Journal of Neural Engineering, 3(3), 189–195. 10.1088/1741-2560/3/3/001

Gonzales, D. L., Khan, H. F., Keri, H. V. S., Yadav, S., Steward, C., Muller, L. E., Pluta, S. R., & Jayant, K. (2025). Touch-evoked traveling waves establish a translaminar spacetime code. Science Advances, 11(5), eadr4038. 10.1126/sciadv.adr4038

Gutierrez, C. A., Lee, C., Kim, B., & Meng, E. (2011). Epoxy-less packaging methods for electrical contact to parylene-based flat flexible cables. 2011 16th International Solid-State Sensors, Actuators and Microsystems Conference, 2299–2302. 10.1109/TRANSDUCERS.2011.5969538

Hahn, N. V., Stein, E., BrainGate Consortium, Donoghue, J. P., Simeral, J. D., Hochberg, L. R., & Willett, F. R. (2025). Long-term performance of intracortical microelectrode arrays in 14 BrainGate clinical trial participants. medRxiv, 2025.07.02.25330310. 10.1101/2025.07.02.25330310

He, F., Lycke, R., Ganji, M., Xie, C., & Luan, L. (2020). Ultraflexible Neural Electrodes for Long-Lasting Intracortical Recording. iScience, 23(8), 101387. 10.1016/j.isci.2020.101387

Jeanpierre, G. M., Omodon, S. C., Goldberg, S. D., Gonzalez, J., Lu, H.-Y., Han, S. S., Baker, M. N., Madariaga, A., Rausch, M. K., Jung, Y., Akinwande, D., Kireev, D., & Santacruz, S. R. (2025). Flexible Microelectrode Arrays with Enhanced Electrochemical Properties Using Platinum Ditelluride. ACS Applied Electronic Materials, acsaelm.5c01581. 10.1021/acsaelm.5c01581

Jun, J. J., Steinmetz, N. A., Siegle, J. H., Denman, D. J., Bauza, M., Barbarits, B., Lee, A. K., Anastassiou, C. A., Andrei, A., Aydın, Ç., Barbic, M., Blanche, T. J., Bonin, V., Couto, J., Dutta, B., Gratiy, S. L., Gutnisky, D. A., Häusser, M., Karsh, B., … Harris, T. D. (2017). Fully integrated silicon probes for high-density recording of neural activity. Nature, 551(7679), 232–236. 10.1038/nature24636

Jung, B., Taylor, P. A., Seidlitz, J., Sponheim, C., Perkins, P., Ungerleider, L. G., Glen, D., & Messinger, A. (2021). A comprehensive macaque fMRI pipeline and hierarchical atlas. NeuroImage, 235, 117997. 10.1016/j.neuroimage.2021.117997

Katsuki and Constantinidis. (2012). Unique and shared roles of the posterior parietal and dorsolateral prefrontal cortex in cognitive functions. Frontiers in Integrative Neuroscience, 6. 10.3389/fnint.2012.00017

Kim, B. J., Kuo, J. T. W., Hara, S. A., Lee, C. D., Yu, L., Gutierrez, C. A., Hoang, T. Q., Pikov, V., & Meng, E. (2013). 3D Parylene sheath neural probe for chronic recordings. Journal of Neural Engineering, 10(4), 045002. 10.1088/1741-2560/10/4/045002

Kozai, T. D. Y., Jaquins-Gerstl, A. S., Vazquez, A. L., Michael, A. C., & Cui, X. T. (2015). Brain Tissue Responses to Neural Implants Impact Signal Sensitivity and Intervention Strategies. ACS Chemical Neuroscience, 6(1), 48–67. 10.1021/cn500256e

Lee, H. C., Gaire, J., Roysam, B., & Otto, K. J. (2018). Placing Sites on the Edge of Planar Silicon Microelectrodes Enhances Chronic Recording Functionality. IEEE Transactions on Biomedical Engineering, 65(6), 1245–1255. 10.1109/TBME.2017.2715811

Lee, K., Paulk, A. C., Ro, Y. G., Cleary, D. R., Tonsfeldt, K. J., Kfir, Y., Pezaris, J. S., Tchoe, Y., Lee, J., Bourhis, A. M., Vatsyayan, R., Martin, J. R., Russman, S. M., Yang, J. C., Baohan, A., Richardson, R. M., Williams, Z. M., Fried, S. I., Hoi Sang, U., … Dayeh, S. A. (2024). Flexible, scalable, high channel count stereo-electrode for recording in the human brain. Nature Communications, 15(1), 218. 10.1038/s41467-023-43727-9

Liu, X., Ren, C., Lu, Y., Liu, Y., Kim, J.-H., Leutgeb, S., Komiyama, T., & Kuzum, D. (2021). Multimodal neural recordings with Neuro-FITM uncover diverse patterns of cortical–hippocampal interactions. Nature Neuroscience, 24(6), Article 6. 10.1038/s41593-021-00841-5

Liu, Y., Jia, H., Sun, H., Jia, S., Yang, Z., Li, A., Jiang, A., Naya, Y., Yang, C., Xue, S., Li, X., Chen, B., Zhu, J., Zhou, C., Li, M., & Duan, X. (2024). A high-density 1,024-channel probe for brain-wide recordings in non-human primates. Nature Neuroscience, 27(8), 1620–1631. 10.1038/s41593-024-01692-6

Lu, Y., Lyu, H., Richardson, A. G., Lucas, T. H., & Kuzum, D. (2016). Flexible Neural Electrode Array Based-on Porous Graphene for Cortical Microstimulation and Sensing. Scientific Reports, 6(1), 33526. 10.1038/srep33526

Luan, L., Wei, X., Zhao, Z., Siegel, J. J., Potnis, O., Tuppen, C. A., Lin, S., Kazmi, S., Fowler, R. A., Holloway, S., Dunn, A. K., Chitwood, R. A., & Xie, C. (2017). Ultraflexible nanoelectronic probes form reliable, glial scar–free neural integration. Science Advances, 3(2), e1601966. 10.1126/sciadv.1601966

Merken, L., Schelles, M., Ceyssens, F., Kraft, M., & Janssen, P. (2022). Thin flexible arrays for long-term multi-electrode recordings in macaque primary visual cortex. Journal of Neural Engineering, 19(6), 066039. 10.1088/1741-2552/ac98e2

Meyer, T., & Constantinidis, C. (2005). A software solution for the control of visual behavioral experimentation. Journal of Neuroscience Methods, 142(1), 27–34. 10.1016/j.jneumeth.2004.07.009

Mozumder, R., Chung, S., Li, S., & Constantinidis, C. (2024). Contributions of narrow- and broad-spiking prefrontal and parietal neurons on working memory tasks. Frontiers in Systems Neuroscience, 18, 1365622. 10.3389/fnsys.2024.1365622

Mozumder, R., & Constantinidis, C. (2023). Single-neuron and population measures of neuronal activity in working memory tasks. Journal of Neurophysiology, 130(3), 694–705. 10.1152/jn.00245.2023

Mozumder, R., Wang, Z., Dang, W., Zhu, J., Hammond, B. M., Machado, A., & Constantinidis, C. (2026). Asynchronous firing and off states in working memory maintenance. Cell Reports, 45(1), 116764. 10.1016/j.celrep.2025.116764

Musk, E., & Neuralink. (2019). An Integrated Brain-Machine Interface Platform With Thousands of Channels. Journal of Medical Internet Research, 21(10), e16194. 10.2196/16194

Namima, T., Kempkes, E., Smith, B., Smith, L., Orsborn, A. L., & Pasupathy, A. (2024). Inserting a Neuropixels probe into awake monkey cortex: Two probes, two methods. Journal of Neuroscience Methods, 402, 110016. 10.1016/j.jneumeth.2023.110016

Oby, E. R., Golub, M. D., Hennig, J. A., Degenhart, A. D., Tyler-Kabara, E. C., Yu, B. M., Chase, S. M., & Batista, A. P. (2019). New neural activity patterns emerge with long-term learning. Proceedings of the National Academy of Sciences, 116(30), 15210–15215. 10.1073/pnas.1820296116

Oh, S., Jekal, J., Won, J., Lim, K. S., Jeon, C.-Y., Park, J., Yeo, H.-G., Kim, Y. G., Lee, Y. H., Ha, L. J., Jung, H. H., Yea, J., Lee, H., Ha, J., Kim, J., Lee, D., Song, S., Son, J., Yu, T. S., … Jang, K.-I. (2025). A stealthy neural recorder for the study of behaviour in primates. Nature Biomedical Engineering, 9(6), 882–895. 10.1038/s41551-024-01280-w

Ortigoza-Diaz, J., Scholten, K., Larson, C., Cobo, A., Hudson, T., Yoo, J., Baldwin, A., Weltman Hirschberg, A., & Meng, E. (2018). Techniques and Considerations in the Microfabrication of Parylene C Microelectromechanical Systems. Micromachines, 9(9), 422. 10.3390/mi9090422

Overton, J. A., Cooke, D. F., Goldring, A. B., Lucero, S. A., Weatherford, C., & Recanzone, G. H. (2017). Improved methods for acrylic-free implants in nonhuman primates for neuroscience research. Journal of Neurophysiology, 118(6), 3252–3270. 10.1152/jn.00191.2017

Pachitariu, M., Rossant, C., & Steinmetz, N. (2021, January 30). MouseLand/Kilosort: Kilosort 2.5. Zenodo. https://zenodo.org/records/4482749

Pachitariu, M., Sridhar, S., Pennington, J., & Stringer, C. (2024). Spike sorting with Kilosort4. Nature Methods, 21(5), 914–921. 10.1038/s41592-024-02232-7

Pachitariu, M., Steinmetz, N. A., Kadir, S. N., Carandini, M., & Harris, K. D. (2016). Fast and accurate spike sorting of high-channel count probes with KiloSort. In D. Lee, M. Sugiyama, U. Luxburg, I. Guyon, & R. Garnett (Eds.), Advances in Neural Information Processing Systems (Vol. 29). Curran Associates, Inc. https://proceedings.neurips.cc/paper_files/paper/2016/file/1145a30ff80745b56fb0cecf65305017-Paper.pdf

Pagan, M., Urban, L. S., Wohl, M. P., & Rust, N. C. (2013). Signals in inferotemporal and perirhinal cortex suggest an untangling of visual target information. Nature Neuroscience, 16(8), 1132–1139. 10.1038/nn.3433

Paulk, A. C., Kfir, Y., Khanna, A. R., Mustroph, M. L., Trautmann, E. M., Soper, D. J., Stavisky, S. D., Welkenhuysen, M., Dutta, B., Shenoy, K. V., Hochberg, L. R., Richardson, R. M., Williams, Z. M., & Cash, S. S. (2022). Large-scale neural recordings with single neuron resolution using Neuropixels probes in human cortex. Nature Neuroscience, 25(2), 252–263. 10.1038/s41593-021-00997-0

Pothof, F., Bonini, L., Lanzilotto, M., Livi, A., Fogassi, L., Orban, G. A., Paul, O., & Ruther, P. (2016). Chronic neural probe for simultaneous recording of single-unit, multi-unit, and local field potential activity from multiple brain sites. Journal of Neural Engineering, 13(4), 046006. 10.1088/1741-2560/13/4/046006

Qi, X., & Constantinidis, C. (2012). Correlated discharges in the primate prefrontal cortex before and after working memory training. European Journal of Neuroscience, 36(11), 3538–3548. 10.1111/j.1460-9568.2012.08267.x

Qiang, Y., Gu, W., Jang, D., Shin, Y., Shi, D., Seo, K. J., Li, G., Vinnikova, S., Wu, S., Iyer, A., Artoni, P., Ryu, J., Bai, T., Dhawan, V., Medalla, M., Rosene, D. L., Moore, T. L., Koppes, A. N., Koppes, R., … Fang, H. (2025). Monolithic three-dimensional neural probes from deterministic rolling of soft electronics. Nature Electronics, 8(8), 721–737. 10.1038/s41928-025-01431-0

Saad, Z. S., Glen, D. R., Chen, G., Beauchamp, M. S., Desai, R., & Cox, R. W. (2009). A new method for improving functional-to-structural MRI alignment using local Pearson correlation. NeuroImage, 44(3), 839–848. 10.1016/j.neuroimage.2008.09.037

Sadtler, P. T., Quick, K. M., Golub, M. D., Chase, S. M., Ryu, S. I., Tyler-Kabara, E. C., Yu, B. M., & Batista, A. P. (2014). Neural constraints on learning. Nature, 512(7515), Article 7515. 10.1038/nature13665

Salatino, J. W., Ludwig, K. A., Kozai, T. D. Y., & Purcell, E. K. (2017). Glial responses to implanted electrodes in the brain. Nature Biomedical Engineering, 1(11), 862–877. 10.1038/s41551-017-0154-1

Santhanam, G., Ryu, S. I., Yu, B. M., Afshar, A., & Shenoy, K. V. (2006). A high-performance brain–computer interface. Nature, 442(7099), Article 7099. 10.1038/nature04968

Song, E., Li, J., Won, S. M., Bai, W., & Rogers, J. A. (2020). Materials for flexible bioelectronic systems as chronic neural interfaces. Nature Materials, 19(6), 590–603. 10.1038/s41563-020-0679-7

Srikantharajah, K., Medinaceli Quintela, R., Doerenkamp, K., Kampa, B. M., Musall, S., Rothermel, M., & Offenhäusser, A. (2021). Minimally-invasive insertion strategy and in vivo evaluation of multi-shank flexible intracortical probes. Scientific Reports, 11(1), 18920. 10.1038/s41598-021-97940-x

Steinmetz, N. A., Zatka-Haas, P., Carandini, M., & Harris, K. D. (2019). Distributed coding of choice, action and engagement across the mouse brain. Nature, 576(7786), 266–273. 10.1038/s41586-019-1787-x

Tang, H., Qi, X.-L., Riley, M. R., & Constantinidis, C. (2019). Working memory capacity is enhanced by distributed prefrontal activation and invariant temporal dynamics. Proceedings of the National Academy of Sciences, 116(14), 7095–7100. 10.1073/pnas.1817278116

Thielen, B., & Meng, E. (2021). A comparison of insertion methods for surgical placement of penetrating neural interfaces. Journal of Neural Engineering, 18(4), 041003. 10.1088/1741-2552/abf6f2

Thielen, B., & Meng, E. (2023). Characterization of thin film Parylene C device curvature and the formation of helices via thermoforming. Journal of Micromechanics and Microengineering, 33(9), 095007. 10.1088/1361-6439/acdc33

Thrower, L., Dang, W., Jaffe, R. G., Sun, J. D., & Constantinidis, C. (2023). Decoding working memory information from neurons with and without persistent activity in the primate prefrontal cortex. Journal of Neurophysiology, 130(6), 1392–1402. 10.1152/jn.00290.2023

Tian, Y., Yin, J., Wang, C., He, Z., Xie, J., Feng, X., Zhou, Y., Ma, T., Xie, Y., Li, X., Yang, T., Ren, C., Li, C., & Zhao, Z. (2023). An Ultraflexible Electrode Array for Large-Scale Chronic Recording in the Nonhuman Primate Brain. Advanced Science, 10(33), 2302333. 10.1002/advs.202302333

Trautmann, E. M., Hesse, J. K., Stine, G. M., Xia, R., Zhu, S., O’Shea, D. J., Karsh, B., Colonell, J., Lanfranchi, F. F., Vyas, S., Zimnik, A., Steinmann, N. A., Wagenaar, D. A., Andrei, A., Lopez, C. M., O’Callaghan, J., Putzeys, J., Raducanu, B. C., Welkenhuysen, M., … Harris, T. (2025). Large-scale high-density brain-wide neural recording in nonhuman primates. Nature Neuroscience, 28, 1562–1575. 10.1038/s41593-025-01976-5

Wang, Q., Yin, J., & Cui, H. (2021). Reinforcement of Neuropixels probes for high-density neural recording in non-human primates. 2021 10th International IEEE/EMBS Conference on Neural Engineering (NER), 128–131. 10.1109/NER49283.2021.9441229

Wang, Y., Wang, Q., Zheng, R., Xu, X., Yang, X., Gui, Q., Yang, X., Wang, Y., Cui, H., & Pei, W. (2023). Flexible multichannel electrodes for acute recording in nonhuman primates. Microsystems & Nanoengineering, 9(1), 93. 10.1038/s41378-023-00550-y

Wang, Z., Singh, B., Zhou, X., & Constantinidis, C. (2022). Strong Gamma Frequency Oscillations in the Adolescent Prefrontal Cortex. The Journal of Neuroscience, 42(14), 2917–2929. 10.1523/JNEUROSCI.1604-21.2022

Wimmer, K., Nykamp, D. Q., Constantinidis, C., & Compte, A. (2014). Bump attractor dynamics in prefrontal cortex explains behavioral precision in spatial working memory. Nature Neuroscience, 17(3), 431–439. 10.1038/nn.3645

Yasar, T. B., Gombkoto, P., Vyssotski, A. L., Vavladeli, A. D., Lewis, C. M., Wu, B., Meienberg, L., Lundegardh, V., Helmchen, F., Von Der Behrens, W., & Yanik, M. F. (2024). Months-long tracking of neuronal ensembles spanning multiple brain areas with Ultra-Flexible Tentacle Electrodes. Nature Communications, 15(1), 4822. 10.1038/s41467-024-49226-9

Zhao, S., Tang, X., Tian, W., Partarrieu, S., Liu, R., Shen, H., Lee, J., Guo, S., Lin, Z., & Liu, J. (2023). Tracking neural activity from the same cells during the entire adult life of mice. Nature Neuroscience, 26(4), 696–710. 10.1038/s41593-023-01267-x

Zhao, Z., Li, X., He, F., Wei, X., Lin, S., & Xie, C. (2019). Parallel, minimally-invasive implantation of ultra-flexible neural electrode arrays. Journal of Neural Engineering, 16(3), 035001. 10.1088/1741-2552/ab05b6

Zhao, Z., Zhu, H., Li, X., Sun, L., He, F., Chung, J. E., Liu, D. F., Frank, L., Luan, L., & Xie, C. (2022). Ultraflexible electrode arrays for months-long high-density electrophysiological mapping of thousands of neurons in rodents. Nature Biomedical Engineering, 7(4), 520–532. 10.1038/s41551-022-00941-y

Zhou, T., Hong, G., Fu, T.-M., Yang, X., Schuhmann, T. G., Viveros, R. D., & Lieber, C. M. (2017). Syringe-injectable mesh electronics integrate seamlessly with minimal chronic immune response in the brain. Proceedings of the National Academy of Sciences, 114(23), 5894–5899. 10.1073/pnas.1705509114

Zhu, J., Garin, C. M., Qi, X.-L., Machado, A., Wang, Z., Ben Hamed, S., Stanford, T. R., Salinas, E., Whitlow, C. T., Anderson, A. W., Zhou, X. M., Calabro, F. J., Luna, B., & Constantinidis, C. (2025). Longitudinal measures of monkey brain structure and activity through adolescence predict cognitive maturation. Nature Neuroscience, 28(11), 2344–2355. 10.1038/s41593-025-02076-0

Zhu, J., Hammond, B. M., Zhou, X. M., & Constantinidis, C. (2023). Laminar pattern of adolescent development changes in working memory neuronal activity. Journal of Neurophysiology, 130(4), 980–989. 10.1152/jn.00294.2023

