## Supplementary Material for "Repeatable, low-drift recordings in behaving non-human primates using flexible microelectrodes"

*Woods, et al.*

Corresponding author:

*Daniel L. Gonzales,*

Document includes:

*Supplementary Discussion*

*Supplementary Figures S1-S6*

### SUPPLEMENTARY DISCUSSION

Unreported studies with flexible arrays

We feel it is important to note that Neuralink has made massive strides in parallel implantation of large-scale, chronic microelectrode arrays in NHPs and humans. While this company provides informal and semi-regular progress updates, their work with NHPs is unpublished and we can only opaquely infer the precise methods and results of their work. Importantly, due to a difference of missions, Neuralink's technological advances are out of reach for the broader scientific community. We see our work as making flexible microelectrode technology accessible to the neuroscience community.

Flexible probe failure modes

The most prominent failure mode was flexible probe breakages. We report 3 probe breakages during the retraction phase of recordings. These failures occurred when inconsistent probe positioning along the microwire—specifically, when the Parylene-C was threaded too far up or seated unevenly—created mechanical stress points during shuttle withdrawal. The exact positioning of the probe relative to the microwire appears critical: too far up and the probe experiences excessive tension during retraction; too shallow and the probe may not remain securely attached during insertion. Other groups have addressed this challenge in rodents (Luan et al., 2017) and NHPs (Tian et al., 2023; Wang et al., 2023) by employing microwire geometries resembling a "thumbtack," where an enlarged head at a fixed position provides a consistent, defined attachment point that securely holds the flexible array tip in place. This design eliminates the variability in probe placement that we encountered and may further improve the implantation yield. However, the thumbtack alone is unlikely to provide reliable retraction and free-floating polymer arrays, as demonstrated in previous NHP studies (Wang et al., 2023).

Prior to implantation, approximately 10% of flexible probes also broke during preparation, specifically while manually threading probes under a microscope. This process is currently laborious compared to traditional rigid microelectrodes and future work should focus on automating or simplifying the threading process, perhaps through custom jigs or pre-assembled probe-shuttle combinations. It should also be noted that some groups have had success implanting in rodents without the need for threading and only using a temporary PEG adhesive between the polymer and microwire (Zhao et al., 2019). In unpublished work, our group performs implantations in this manner in mice as well; however, we have found threading to be critical for successful NHP implantations.

Electrode array design

Our investigation shows the first quantitative evidence that metal contact position along a thin polymer array affects recording quality, specifically single-unit yield. We made a data-driven decision to transition from our original checkerboard array design to a linear design based on both probe yield and electrode performance (Figure S4). Evidence of these geometric constraints has also been seen for silicon shanks (Fiáth et al., 2021; H. C. Lee et al., 2018) but never replicated for flexible arrays. We believe that edge electrodes have enhanced tissue contact and decreased distance to nearby neurons, improving the likelihood of single-unit recordings. This does not necessarily mean improved SNR or spike amplitude (Fig. S3-4). This geometric constraint may be the reason that other polymer arrays with electrodes placed near the center of wide, flexible shanks reported a low yield of single-unit activity during NHP recordings (Jeanpierre et al., 2025; K. Lee et al., 2024; Oh et al., 2025). These findings should strongly inform future design decisions for researchers developing neural interfaces.

Low drift Si recordings

Our flexible arrays in large animal models displayed dramatic reductions in single-unit drift. It should be noted that low-drift recordings with rigid arrays are possible. A recent study using Neuropixels 1.0 NHP reported a low drift similar to those we should here with flexible electrodes (Trautmann et al., 2025). However, this stability also requires long “settling” periods after insertion lasting 30-60 min prior to recording, which is not always suitable for maintaining animal motivation during behavioral tasks. In addition, this study also used a blunt guide tube pressed against the brain surface to minimize motion and swelling. Our flexible electrodes achieve superior stability without these constraints.

### SUPPLEMENTAL FIGURES

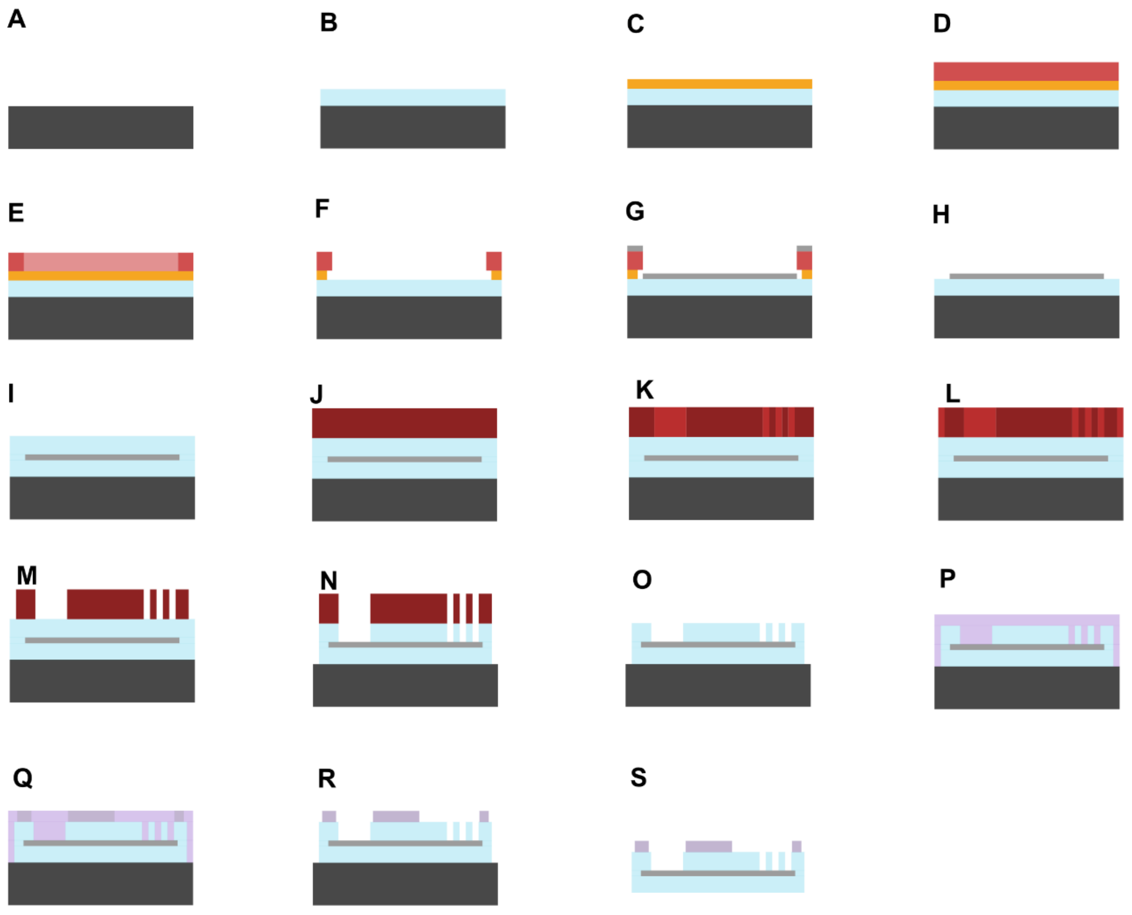


**Figure S1: Full fabrication of flexible microelectrode arrays**

(A) Si wafer substrate, (B) Parylene-C CVD, (C) LOR7A, (D) S1813, (E) Metal layer UV exposure, and (F) developing in MF-319, (G) Pt sputter deposition, (H) liftoff in remover PG, (I) 2$^{nd}$ Parylene-C CVD, (J) SPR 7.0, (K) contact and connection pad UV exposure, (L) probe outline UV exposure, (M) probe electrode and outline development, (N) probe electrode and outline etching, (O) strip SPR 7.0, (P) coat in SU-8, (Q) UV expose probe tip and backend, (R) develop SU-8, (S) peel probe from Si substrate

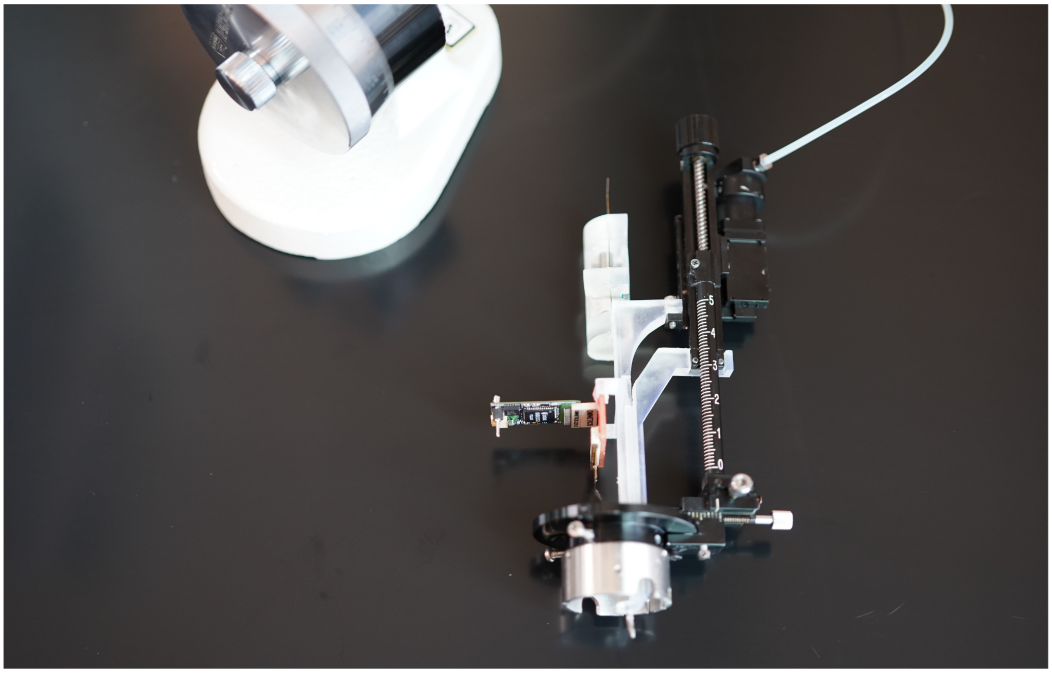

**Figure S2: Microdrive adapter**

Image shows Narishige MO-97A microdrive with our custom 3D printed drive adapter holding a loaded probe with the custom PCB and Intan headstage.

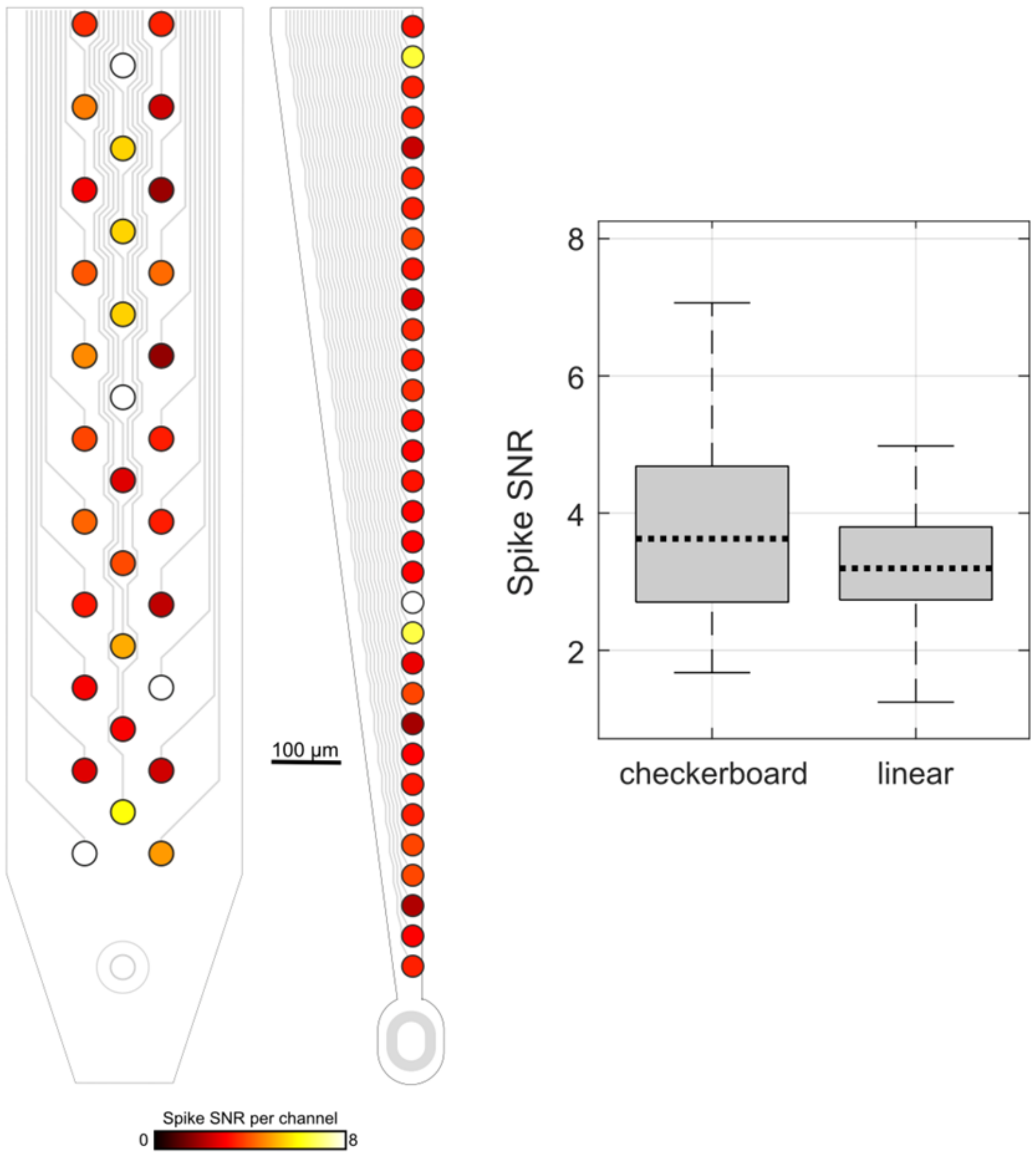


**Figure S3: Signal to Noise ratio between probe types**

(Left and Center) Spike SNR per channel for both flexible array geometries, (Right) Median spike SNR across channel by probe type.

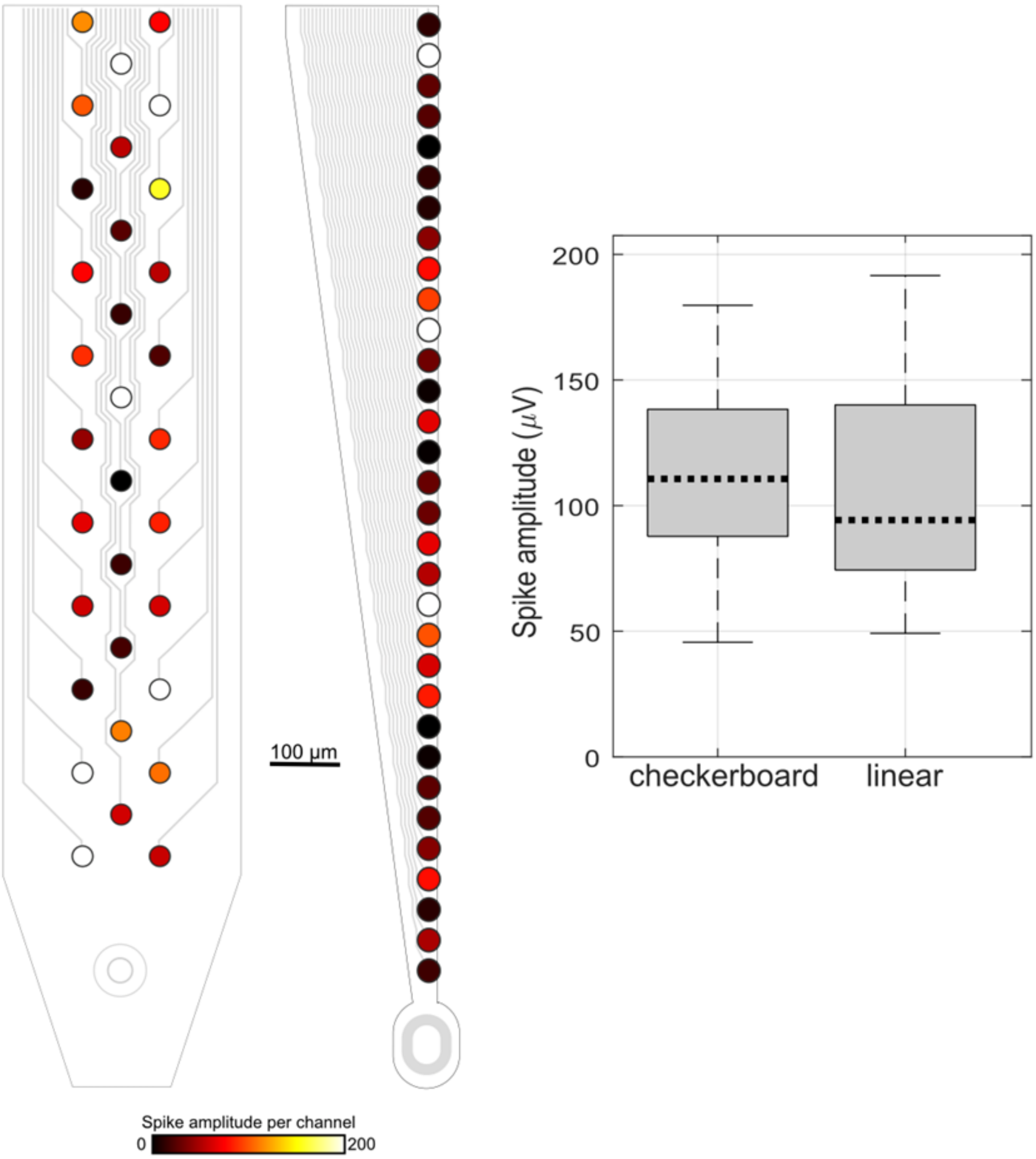


**Figure S4: Unit amplitude between probe types**

(Left and Center) Spike amplitude per channel for both flexible array geometries, (Right) Median spike amplitude across channels by probe type.

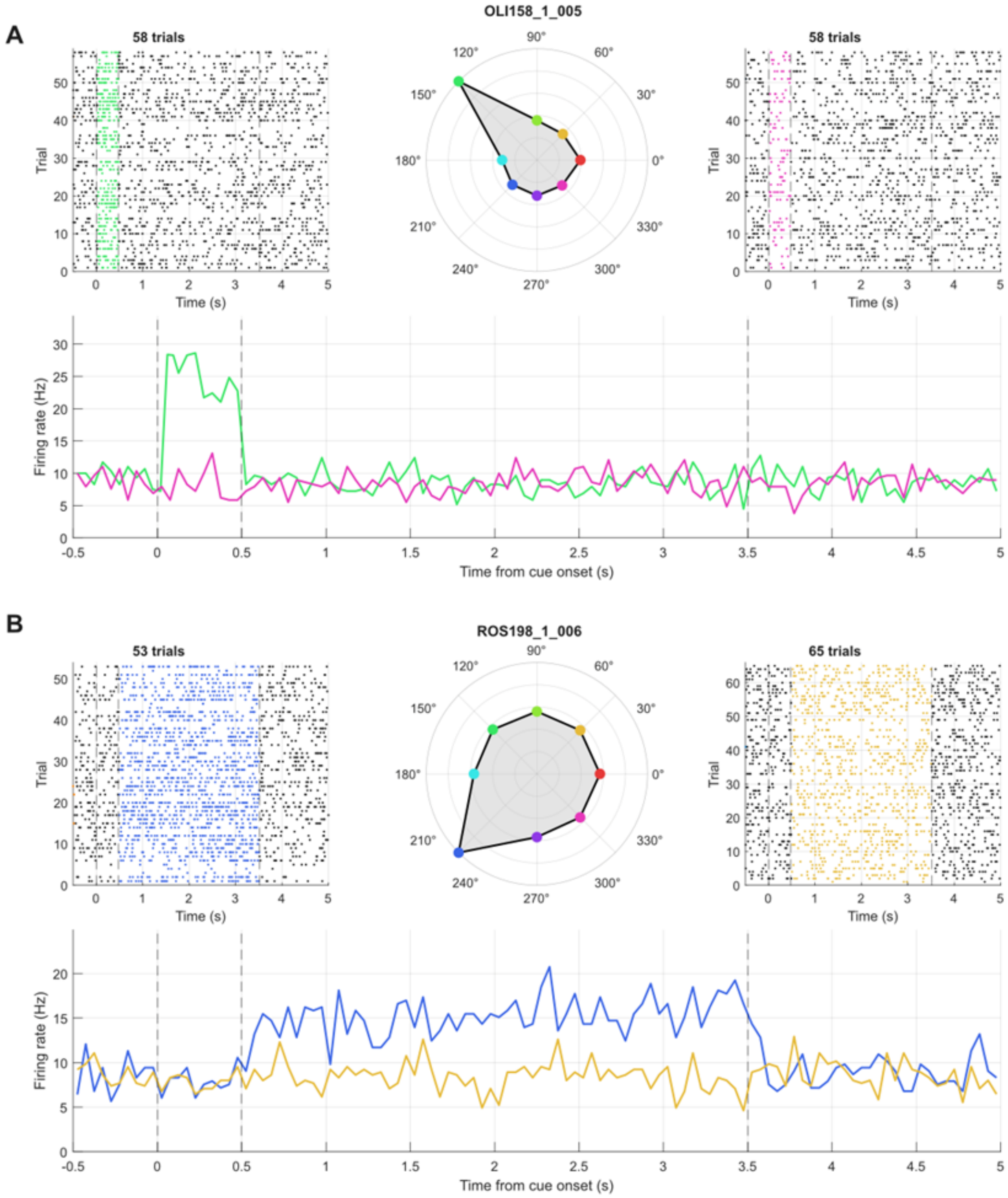


**Figure S5: Representative cue and delay cells**

**(A)** The same cells are shown in Fig. 5B. (Top left and right) Raster plots showing the preferred and diametric cue location for selected unit. (Top center) Polar plot showing preferred cue location for this cell. (Bottom) PSTH showing preferred (green) and opposite (cyan) traces for this cue responsive cell

**(B)** (Top left and right) Raster plots showing the preferred and diametric cue location for selected unit. (Top center) Polar plot showing preferred cue location for this cell. (Bottom) PSTH showing preferred (blue) and diametric (yellow) traces for this delay responsive cell

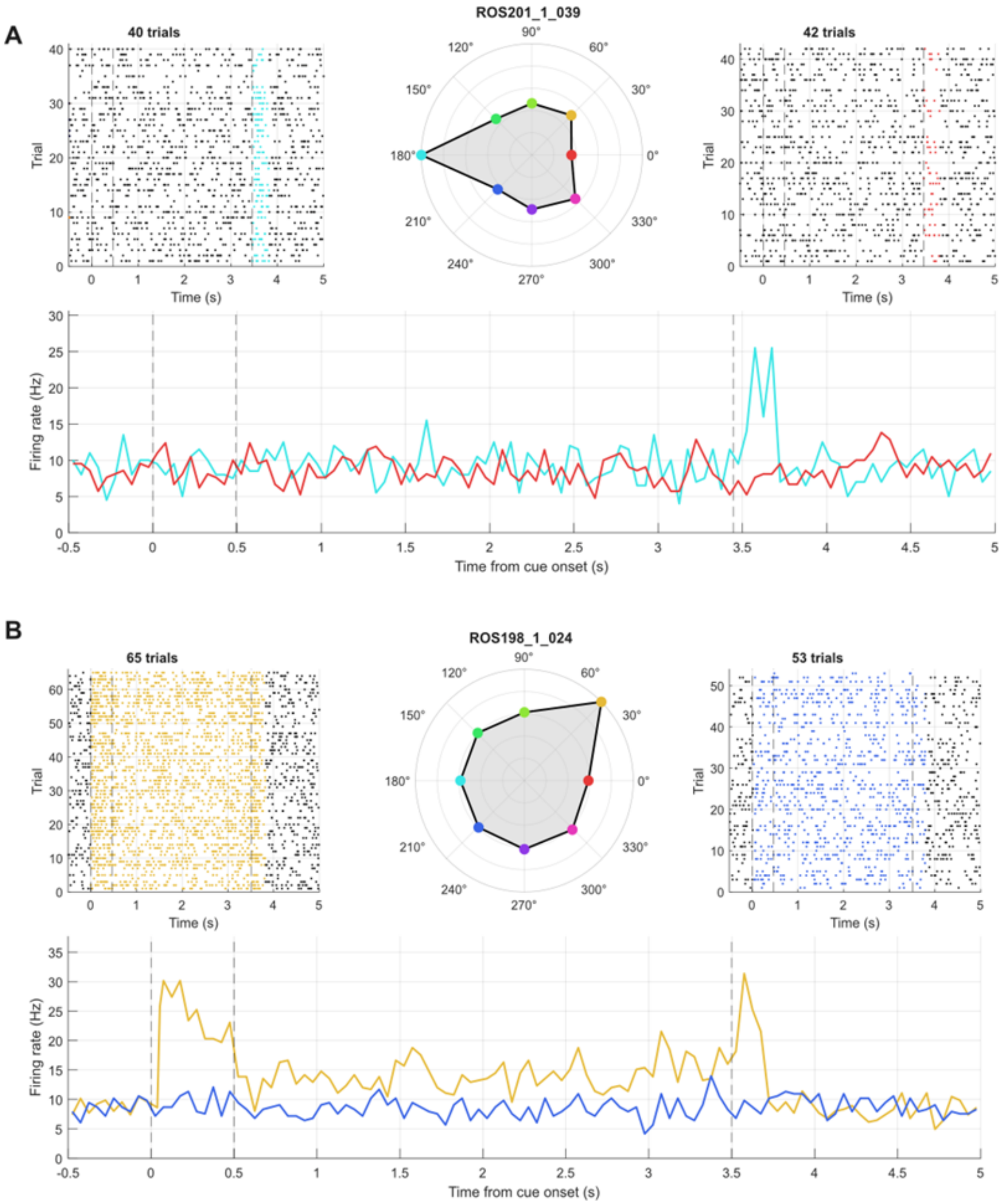


**Figure S6: Representative saccade and multiple epoch cells**

**(A)** (Top left and right) Raster plots showing the preferred and diametric cue location for selected unit. (Top center) Polar plot showing preferred cue location for this cell. (Bottom) PSTH showing preferred (blue) and diametric (red) traces for this saccade responsive cell

**(B)** (Top left and right) Raster plots showing the preferred and diametric cue location for selected unit. (Top center) Polar plot showing preferred cue location for this cell. (Bottom) PSTH showing preferred (yellow) and diametric (blue) traces for this multi-epoch responsive cell

### SUPPLEMENTARY REFERENCES


Fiáth, R., Meszéna, D., Somogyvári, Z., Boda, M., Barthó, P., Ruther, P., & Ulbert, I. (2021). Recording site placement on planar silicon-based probes affects signal quality in acute neuronal recordings. *Scientific Reports*, *11*(1), 2028. https://doi.org/10.1038/s41598-021-81127-5

Jeanpierre, G. M., Omodon, S. C., Goldberg, S. D., Gonzalez, J., Lu, H.-Y., Han, S. S., Baker, M. N., Madariaga, A., Rausch, M. K., Jung, Y., Akinwande, D., Kireev, D., & Santacruz, S. R. (2025). Flexible Microelectrode Arrays with Enhanced Electrochemical Properties Using Platinum Ditelluride. *ACS Applied Electronic Materials*, acsaelm.5c01581. https://doi.org/10.1021/acsaelm.5c01581

Lee, H. C., Gaire, J., Roysam, B., & Otto, K. J. (2018). Placing Sites on the Edge of Planar Silicon Microelectrodes Enhances Chronic Recording Functionality. *IEEE Transactions on Biomedical Engineering*, *65*(6), 1245–1255. https://doi.org/10.1109/TBME.2017.2715811

Lee, K., Paulk, A. C., Ro, Y. G., Cleary, D. R., Tonsfeldt, K. J., Kfir, Y., Pezaris, J. S., Tchoe, Y., Lee, J., Bourhis, A. M., Vatsyayan, R., Martin, J. R., Russman, S. M., Yang, J. C., Baohan, A., Richardson, R. M., Williams, Z. M., Fried, S. I., Hoi Sang, U., … Dayeh, S. A. (2024). Flexible, scalable, high channel count stereo-electrode for recording in the human brain. *Nature Communications*, *15*(1), 218. https://doi.org/10.1038/s41467-023-43727-9

Luan, L., Wei, X., Zhao, Z., Siegel, J. J., Potnis, O., Tuppen, C. A., Lin, S., Kazmi, S., Fowler, R. A., Holloway, S., Dunn, A. K., Chitwood, R. A., & Xie, C. (2017). Ultraflexible nanoelectronic probes form reliable, glial scar–free neural integration. *Science Advances*, *3*(2), e1601966. https://doi.org/10.1126/sciadv.1601966

Oh, S., Jekal, J., Won, J., Lim, K. S., Jeon, C.-Y., Park, J., Yeo, H.-G., Kim, Y. G., Lee, Y. H., Ha, L. J., Jung, H. H., Yea, J., Lee, H., Ha, J., Kim, J., Lee, D., Song, S., Son, J., Yu, T. S., … Jang, K.-I. (2025). A stealthy neural recorder for the study of behaviour in primates. *Nature Biomedical Engineering*, *9*(6), 882–895. https://doi.org/10.1038/s41551-024-01280-w

Tian, Y., Yin, J., Wang, C., He, Z., Xie, J., Feng, X., Zhou, Y., Ma, T., Xie, Y., Li, X., Yang, T., Ren, C., Li, C., & Zhao, Z. (2023). An Ultraflexible Electrode Array for Large-Scale Chronic Recording in the Nonhuman Primate Brain. *Advanced Science*, *10*(33), 2302333. https://doi.org/10.1002/advs.202302333

Trautmann, E. M., Hesse, J. K., Stine, G. M., Xia, R., Zhu, S., O'Shea, D. J., Karsh, B., Colonell, J., Lanfranchi, F. F., Vyas, S., Zimnik, A., Steinmann, N. A., Wagenaar, D. A., Andrei, A., Lopez, C. M., O'Callaghan, J., Putzeys, J., Raducanu, B. C., Welkenhuysen, M., … Harris, T. (2025). Large-scale high-density brain-wide neural recording in nonhuman primates. *Nature Neuroscience*, *28*, 1562–1575. https://doi.org/10.1038/s41593-025-01976-5

Wang, Y., Wang, Q., Zheng, R., Xu, X., Yang, X., Gui, Q., Yang, X., Wang, Y., Cui, H., & Pei, W. (2023). Flexible multichannel electrodes for acute recording in nonhuman primates. *Microsystems & Nanoengineering*, *9*(1), 93. https://doi.org/10.1038/s41378-023-00550-y

Zhao, Z., Li, X., He, F., Wei, X., Lin, S., & Xie, C. (2019). Parallel, minimally-invasive implantation of ultra-flexible neural electrode arrays. *Journal of Neural Engineering*, *16*(3), 035001. https://doi.org/10.1088/1741-2552/ab05b6
